# Feedback modulation of Orai1α and Orai1β protein content mediated by STIM proteins

**DOI:** 10.1101/2024.03.05.583469

**Authors:** Joel Nieto-Felipe, Alvaro Macias-Díaz, Vanesa Jimenez-Velarde, Jose J. Lopez, Gines M. Salido, Tarik Smani, Isaac Jardin, Juan A. Rosado

## Abstract

Store-operated Ca^2+^ entry (SOCE) is a mechanism controlled by the filling state of the intracellular Ca^2+^ stores, predominantly the endoplasmic reticulum (ER), where ER-resident proteins STIM1 and STIM2 orchestrate the activation of Orai channels in the plasma membrane, and Orai1 playing a predominant role. Two forms of Orai1, Orai1α and Orai1β, have been identified, which arises the question whether they are equally regulated by STIM proteins. We demonstrate that STIM1 preferentially activates Orai1α over STIM2, yet both STIM proteins similarly activate Orai1β. Under resting conditions, there is a pronounced association between STIM2 and Orai1α, implying their significance in maintaining basal Ca^2+^ levels within the cytosol and ER. STIM1 and STIM2 are also shown to influence the protein levels of the Orai1 variants, independent of Ca^2+^ influx, via lysosomal degradation. Interestingly, Orai1α and Orai1β appear to selectively regulate the protein level of STIM1, but not STIM2. These observations offer crucial insights into the regulatory dynamics between STIM proteins and Orai1 variants, enhancing our understanding of the intricate processes that fine-tune intracellular Ca^2+^ signaling.

## Introduction

The tetra-pass plasma membrane (PM) protein Orai1 belongs to a highly selective Ca^2+^ channel family, ubiquitously conserved throughout the animal kingdom, that are known for their vital role in the finely modulated spatio-temporal signaling of Ca^2+^ across the PM. Working in conjunction with its mammalian paralogues, Orai2 and Orai3, Orai1 is instrumental in facilitating store-operated calcium entry (SOCE), a fundamental process essential for agonist-induced Ca^2+^ mobilization. This mobilization regulates various cellular functions, including gene expression, cell proliferation, differentiation, and the immune response in both electrically excitable and non-excitable cells [1-3].

SOCE is initiated by the release of Ca^2+^ from intracellular stores, primarily the endoplasmic reticulum (ER). This reduction in the luminal Ca^2+^ concentration is detected by stromal interaction molecule proteins, STIM1 and STIM2, a type of integral single-pass ER proteins that possess highly conserved luminal EF-hand motifs, which have the ability to bind Ca^2+^, enabling STIM proteins to effectively detect and monitor stored Ca^2+^ levels within the ER [4]. STIM1 and STIM2 exhibit distinct characteristics and biological roles. STIM1 primarily functions as the principal stimulator of Orai/CRAC channels, whereas STIM2, with less affinity for Ca^2+^ ions, acts as a comparatively slower and less potent activator of Orai channels, and it is considered to be involved in maintaining stable levels of Ca^2+^ in both, the cytosol and ER, under resting conditions [5]. Upon ER depletion, STIM proteins activate, oligomerize, and translocate to regions close to the PM. There, they associate with and initiate the activation of Orai channels through a specific cytosolic region located within the STIM C-terminus [6-8]. This region, known as Orai1 activating small fragment (OASF) [6], STIM1-Orai1 activating region (SOAR) [7] or CRAC activation domain (CAD) [8], includes two coiled-coil structures, highly conserved among family members, containing a homomerization domain, required for coupling to Orai1, and a regulatory domain, which modulates the extent of coupling to Orai1 [6]. Subsequently, the activation of Orai channels leads to a significant influx of Ca^2+^ from the extracellular environment [9, 10].

Currently, there are no reported cases of alternative splicing for Orai1 mRNA; however, alternative initiation of translation has been observed in mammalian cells, giving rise to two Orai1 isoforms. The canonical full-length Orai1 protein, designated as Orai1α[, comprises 301 amino acids, whereas a shorter variant, termed Orai1[β, originates from an alternative initiation of translation commencing at either methionine 64 or methionine 71. This variant consequently lacks the N-terminal 63 or 70 amino acids found in the full-length Orai1 protein [11, 12]. While both variants of Orai1 demonstrate similar efficacy in supporting the highly Ca^2+^-selective and store-operated *I_CRAC_* currents [13], notable functional and biophysical distinctions exist between Orai1α and Orai1β. Functionally, Orai1α, in conjunction with STIM1 and TRPC1, mediates the less Ca^2+^-selective store-operated current *I_SOC_* [14-16] and the arachidonate-regulated current *I_ARC_* [13]. In contrast, Orai1β does not participate in *I_ARC_*, and its involvement in *I_SOC_*is cell-type-dependent [17]. Furthermore, Orai1α is essential for the NF-κB transcription factor activation, while Orai1β does not exhibit this capability [18]. Biophysically, Orai1α has been shown to be more susceptible to rapid Ca^2+^-dependent inactivation compared to Orai1β [13]. However, the interaction patterns between STIM1 or STIM2 and Orai1 variants, as well as their effectiveness in activating Orai1α and Orai1β, have not been studied to date.

In a resting cell, Orai1 channels are present in two distinct cellular regions, the PM, which accounts for approximately 40%, and a sub-plasmalemal compartment [19]. Upon cell stimulation, intracellular Ca^2+^ store depletion leads to an accumulation of Orai1 proteins in the PM, in a STIM1-dependent mechanism known as Orai1 “trafficking trap” that reduces Orai1 recycling and compartmentalization, thereby supporting and sustaining SOCE [19]. As we have previously reported, this mechanism requires the participation of SNAP-25 [20]. We have recently demonstrated that while both Orai1 variants constitutively translocates to the PM in resting cells, Orai1[β required of Orai1α to adequately translocate to the PM upon store depletion and support SOCE [21]. Furthermore, Orai1 undergoes rapid and dynamic recycling at the PM via endocytosis, process modulated by the chaperonin[containing T[complex protein 1 (CCT) [22]. However, it is still unknown whether both variants of Orai1 are subjected to the same mechanism.

The objective of this study is to investigate the effectiveness of STIM1 and STIM2 in activating Ca^2+^ signals through Orai1α and Orai1β and to characterize the role of STIM proteins in the turnover of Orai1α and Orai1β. Here we report two important aspects of the regulation of Orai1 variants. Firstly, while STIM1 is a stronger activator of Orai1α compared to STIM2, both isoforms activate Orai1β-dependent Ca^2+^ signals to a similar extent. Additionally, we observed a higher basal interaction between STIM2 and Orai1α, suggesting that Orai1α may play a significant role in regulating basal Ca^2+^ levels in the cytosol and ER. Secondly, our results indicate that the expression of STIM proteins can modulate the protein content of Orai1α and Orai1β. This suggests that the regulation of SOCE may involve a feedback mechanism where the extent of Ca^2+^ influx through these channels depends not only on their intrinsic properties but also on the number and location of available channels in the PM. Therefore, our study provides valuable insights into the activation of Orai1α and Orai1β by STIM1 and STIM2, as well as the differential regulation of their expression. These findings contribute to a better understanding of the complex mechanisms underlying Ca^2+^ signaling and the role of Orai1 variants in cellular functions.

## Materials and methods

### Reagents and antibodies

Fura-2 acetoxymethyl ester (fura-2/AM) was from Molecular Probes (Leiden, The Netherlands). High-glucose Dulbecco’s modified Eagle’s medium (DMEM), fetal bovine serum, trypsin, penicillin/streptomycin, Trizma base, streptavidin-conjugated agarose beads, rabbit polyclonal anti-STIM2 antibody (catalog number STIM2-201AP, epitope: amino acids 600-650 of human STIM2), mouse monoclonal anti-PMCA antibody (clone 5F10; catalog number MA3-914, epitope: amino acids 724–783 of human PMCA), Clean-Blot™ IP detection reagent, SuperSignal® West Dura extended duration substrate reagent, sulfo-NHS-LC biotin, Live/Dead viability/cytotoxicity kit and Pierce™ BCA protein assay kit were purchased from ThermoFisher Scientific (Waltham, MA, USA). Complete EDTA-free protease inhibitor cocktail tablets were from Roche Diagnostics GmbH (Mannheim, Germany). DharmaFECT kb transfection reagent was obtained from Cultek (Madrid, Spain). Thapsigargin (TG), carbachol (CCh), protein A agarose beads, cycloheximide, HEPES (4-(2-Hydroxyethyl)piperazine-1-ethanesulfonic acid), EGTA (ethylene glycol-bis(2-aminoethylether)-N,N,N′,N′-tetraacetic acid), EDTA (ethylenedinitrilotetraacetic acid), bovine serum albumin (BSA), sodium azide, dimethyl-BAPTA, sodium ascorbate, bafilomycin A1, MG132, rabbit polyclonal anti-Orai1 antibody (catalog number O8264, epitope: amino acids 288-301 of human Orai1) and rabbit polyclonal anti-β-actin antibody (catalog number A2066, epitope: amino acids 365-375 of human β-actin) were obtained from Sigma (Madrid, Spain). Mouse monoclonal Anti-GOK/STIM1 antibody (Clone 44/GOK; catalog number 610954, epitope: amino acids: 25–139 of human STIM1) was purchased from BD Biosciences (San Jose, CA, USA). Horseradish peroxidase-conjugated goat anti-mouse immunoglobulin G (IgG) antibody and goat anti-rabbit IgG antibody were from Jackson laboratories (West Grove, PA, USA). Thymidine kinase (TK)- and CMV-promoter Orai1α-eGFP and Orai1β-eGFP plasmids were kindly provided by Mohamed Trebak (Department of Pharmacology and Chemical Biology, University of Pittsburgh, Pittsburgh, PA, USA). CMV-promoter STIM1-D76A, STIM1-YFP, STIM2-YFP, STIM1-mCherry and STIM2-mCherry were a gift from Christoph Romanin (Institute of Biophysics, Johannes Kepler University Linz, Linz, Austria). pEX-CMV-SP-YFP-STIM2(15-746) EF hand D80A was a gift from Tobias Meyer (Addgene plasmid # 18864; http://n2t.net/addgene:18864; RRID:Addgene_18864). N-glycosidase F (PNGaseF) from *Elizabethkingia miricola* was from New England Biolabs (Ipswich, MA, USA). All other reagents were of an analytical grade.

### Cell culture and transfections

CRISPR-generated STIM1/2 double-knockout HEK293 cells (DKO) and parental HEK-293 cells were kindly provided by Mohamed Trebak and cultured at 37 °C with a 5% CO_2_ in high-glucose DMEM supplemented with 10% (v/v) fetal bovine serum and 100 U/mL penicillin and streptomycin, as described previously [23]. Cells used throughout the study are free of mycoplasma. For transient transfections, cells were grown to 60 to 80% confluency and transfected with TK- or CMV-promoter Orai1α-EGFP and Orai1β-EGFP or CMV-promoter STIM1-YFP, STIM2-YFP or STIM2-D80A-YFP, depending on the experimental conditions, using DharmaFECT kb transfection reagent and were used 48 h after transfection. Unless otherwise stated, expression plasmids were used at the Orai1:STIM ratio of 1:2 to maintain Orai1-STIM stoichiometry as previously established [24]. Cell viability after transfection throughout the study, estimated using the Live/Dead viability/cytotoxicity kit, was in the range between 92 and 95%. For Western blotting and immunoprecipitation, cells (2 × 10^6^) were plated in 100-mm petri dish and cultured for 48 h, while, for Ca^2+^ imaging and confocal analysis cells (4 × 10^5^) were seeded in a 35-mm six-well multidish.

### Immunoprecipitation and Western blotting

Immunoprecipitation and Western blotting were performed as described previously [25]. Briefly, cells cultured on 100-mm petri dish (8 × 10^6^ cells) were stimulated with 2 μM TG or with vehicle and subsequently lysed with ice-cold NP-40 buffer pH 8 containing 137 mM NaCl, 20 mM Tris, 2 mM EDTA, 10% glycerol, 1% Nonidet P-40, 1 mM Na_3_VO_4_, and complete EDTA-free protease inhibitor tablets. Cell lysates (1 mL) were immunoprecipitated by incubation with 2 µg of anti-Orai1 antibody and 50 µL of protein A-agarose overnight at 4 °C on a rotary platform. Cell lysates and immunoprecipitates were resolved by 10% SDS-PAGE and separated proteins were electrophoretically transferred onto nitrocellulose membranes for subsequent probing. Blots were incubated overnight with 10% (w/v) BSA in Tris-buffered saline with 0.1% Tween-20 (TBST) to block residual protein binding sites. Immunodetection of STIM1, STIM2, Orai1 or β-actin was achieved by incubation for 1h with anti-STIM1 antibody diluted 1:500 in TBST, 1h with anti-STIM2 antibody diluted 1:1000 in TBST, 1 h with anti-Orai1 antibody diluted 1:1000 in TBST or 45 min with anti-β-actin antibody diluted 1:2000 in TBST. The primary antibody was removed, and blots were washed six times for 5 min each with TBST. To detect the primary antibody, blots were incubated for 1 h with horseradish peroxidase-conjugated goat anti-mouse IgG antibody, horseradish peroxidase-conjugated goat anti-rabbit IgG antibody diluted 1:10000 in TBST, or Clean-Blot™ IP Detection Reagent diluted 1:250 in TBST, and then exposed to enhanced chemiluminiscence reagents for 5 min. The antibody binding was assessed with a ChemiDoc Imaging System (Bio-Rad, Madrid, Spain) and the density of bands was measured using ImageJ software v.1.8.0_172 (NIH, Bethesda, MD, USA). Data were normalized to the amount of protein recovered by the antibody used for the immunoprecipitation or to β-actin from the same gel.

### Determination of cytosolic free-Ca^2+^ concentration ([Ca^2+^]_i_)

Cells were loaded with fura-2 by incubation with 5 μM fura-2/AM for 30 min at 37 °C. Coverslips with cultured cells were mounted on a perfusion chamber and placed on the stage of an epifluorescence inverted microscope (Nikon Eclipse Ti2, Amsterdam, The Netherlands) with an image acquisition and analysis system for videomicroscopy (NIS-Elements Imaging Software v.5.02.00, Nikon, Amsterdam, The Netherlands). Cells were continuously superfused at room temperature with HEPES-buffered saline (HBS) containing (in mM) 125 NaCl, 5 KCl, 1 MgCl_2_, 5 glucose, and 25 HEPES, pH 7.4, supplemented with 0.1% (*w*/*v*) BSA. Cells were examined at 40× magnification (Nikon CFI S FLUOR 40× Oil, Amsterdam, The Netherlands) and were alternatively excited with light from a xenon lamp passed through a high-speed monochromator Optoscan ELE 450 (Cairn Research, Faversham, UK) at 340/380 nm. Fluorescence emission at 510 nm was detected using a cooled digital sCMOS camera PCO Panda 4.2 (Excelitas PCO GmbH, Germany) and recorded using NIS-Elements AR software (Nikon, Amsterdam, The Netherlands). Fluorescence ratio (F340/F380) was calculated pixel by pixel, and the data were presented as ΔF_340_/F_380_ as described previously [17]. CCh-evoked Ca^2+^ mobilization was estimated as the area under the curve measured as the integral of the rise in fura-2 fluorescence ratio 10 min after the addition of the agonist and taking a sample every second. TG-evoked Ca^2+^ entry was estimated as the area under the curve measured as the integral of the rise in fura-2 fluorescence ratio 2½ min after the addition of TG and taking a sample every second.

### Analysis of Ca^2+^ oscillations

The analysis of Ca^2+^ oscillations was performed as described previously [26]. All Ca^2+^ traces obtained in the Ca^2+^-imaging experiments were plotted using GraphPad Prism v.8.4.3 (GraphPad Software, San Diego, CA, USA). The number of oscillations per 20 min were counted by the function signal.find_peaks from the SciPy 1.8.1 library for Python 3.9. Then, cells were classified in two groups and a percentage of each group of cells was calculated for each individual coverslip. The first group, oscillating cells, includes cells showing regenerative oscillations after CCh stimulation, where each oscillation returns to baseline before the start of the next oscillation. The second group, non-oscillating cells, includes cells showing a sustained or transient cytosolic Ca^2+^ signal for at least 3 min after stimulation.

### Confocal microscopy analysis

Cells were seeded in 35-mm six-well multidish and transfected with the indicated plasmids. To determine subcellular location of proteins and Förster resonance energy transfer (FRET) cells were imaged 48 h post transfection using a confocal microscope (LSM900, Zeiss, Oberkochen, Germany) with 63× oil immersion objective, using an image acquisition and analysis system for video microscopy (ZEN Software, Zeiss, Oberkochen, Germany). For FRET analysis, appropriate cross-talk calibration factors were determined for each construct on each raw of FRET experiments. After threshold determination and background subtraction, the corrected FRET (Eapp) was calculated on a pixel-to-pixel basis with a custom-made software integrated in FIJI ImageJ 1.54f according to the method published previously [27]. To assess the expression of Orai1 variants in the presence or absence of STIM proteins, background subtraction and thresholding techniques were applied. The NaN Background mathematical process was used to eliminate noise. Subsequently, seven to ten random locations, measuring 2.5 µm^2^ each, were selected for fluorescence measurement using custom software integrated into FIJI ImageJ 1.54f.

### Biotinylation of surface proteins

Biotinylation of surface proteins was performed as described previously [21]. Briefly, cells were washed three times with phosphate-buffered saline (PBS, 137 mM NaCl, 2.7 mM KCl, 1.5 mM KH_2_PO_4_, 8 mM Na_2_HPO_4_•2H_2_O, pH 7,4), subsequently resuspended in biotinylation buffer (PBS, pH 8.0) and surface labeled with 1 mg/mL sulfo-NHS-LC biotin at room temperature. Labeling was stopped 1 h after reaction with PBS, pH 8.0, supplemented with 100 µM Tris and washed 2 times in supplemented PBS, pH 8.0. Biotinylated cells were subsequently lysed with ice-cold NP-40 buffer, pH 8.0, containing 137 mM of NaCl, 20 mM of Tris, 2 mM of EDTA, 10% glycerol, 1% Nonidet P-40, 1 mM of Na_3_VO_4_, and complete EDTA-free protease inhibitor tablets and protein lysates were incubated with streptavidin-conjugated agarose beads overnight at 4 °C on a rocking platform. Biotinylated proteins bound to streptavidin-conjugated agarose beads were isolated by centrifugation and washed 3 times in NP40 buffer. Biotinylated fraction was loaded and separated in 10% sodium dodecyl sulfate-polyacrylamide gel electrophoresis (SDS-PAGE) and analyzed by Western blot using anti-PMCA antibody, as control.

### Statistical analysis

All data are presented as the mean ± standard error of mean (SEM). Analysis of statistical significance was performed using GraphPad Prism v.8.4.3 (GraphPad Software, San Diego, CA, USA). Kruskal–Wallis test combined with Dunńs post hoc test were used to compare the different experimental groups. For comparison between two groups, the Mann–Whitney U test was used. All data with *p* < 0.05 was deemed significant; “ns”[=[nonsignificant.

## Results

### Efficacy of STIM1 and STIM2 activating Ca^2+^ signals through Orai1α and Orai1β

To investigate the functional interaction of Orai1 variants and STIM proteins we have explored thapsigargin (TG)-induced SOCE in STIM1/2 double-knockout HEK-293 cells (DKO cells) co-transfected with CMV-driven STIM1-YFP or STIM2-YFP together with Orai1α-eGFP or Orai1β-eGFP. The expression plasmids were used at the Orai1:STIM ratio of 1:2 to maintain Orai1-STIM stoichiometry as previously reported [28]. Forty-eight hours after transfection of STIM1-YFP or STIM2-YFP, a single protein band with molecular weight around 105 and 130 kDa were observed (Fig. 1F). Furthermore, 48 h after transfection of Orai1α-eGFP or Orai1β-eGFP several bands over 60 and 50 kDa, respectively, were observed, due to Orai1α and Orai1β glycosylation, as previously reported [11, 17]. In the absence of extracellular Ca^2+^, cell treatment with 2 µM TG, an inhibitor of the sarco/endoplasmic reticulum Ca^2+^-ATPase (SERCA), induced a transient increase in the fura-2 fluorescence ratio due to Ca^2+^ release from the intracellular Ca^2+^ stores. Subsequent addition of Ca^2+^ to the extracellular medium leads to a rise in the fura-2 fluorescence ratio that is indicative of Ca^2+^ influx (Fig. 1A). As shown in Fig. 1F, DKO cells do not express STIM proteins and show impaired SOCE (Fig. 1A) despite Ca^2+^ release from the intracellular stores was similar to that in WT cells. Co-transfection with Orai1α or Orai1β with STIM1 rescued SOCE in DKO cells (Figs. 1B-E) resulting in an extent of Ca^2+^ influx that exceeded that in WT cells (probably due to the overexpression of the Orai1 variants and STIM1); however, in DKO cells, co-transfection of STIM2 with Orai1α was less effective than STIM1 (Figs. 1B, D and E). Meanwhile, the efficiency of STIM2 stimulating Ca^2+^ entry through Orai1β was similar to that of STIM1 (Figs. 1C-E). Finally, co-expression of STIM1 and STIM2 with the Orai1 variants generated SOCE with similar extent as STIM1 alone (Figs. 1B-E). These findings indicate that STIM2 is less effective inducing SOCE by Orai1α but exhibit a similar efficacy to STIM1 activating Orai1β. Fig. 1F shows the expression of the target proteins under the different experimental conditions.

**Fig. 1.**
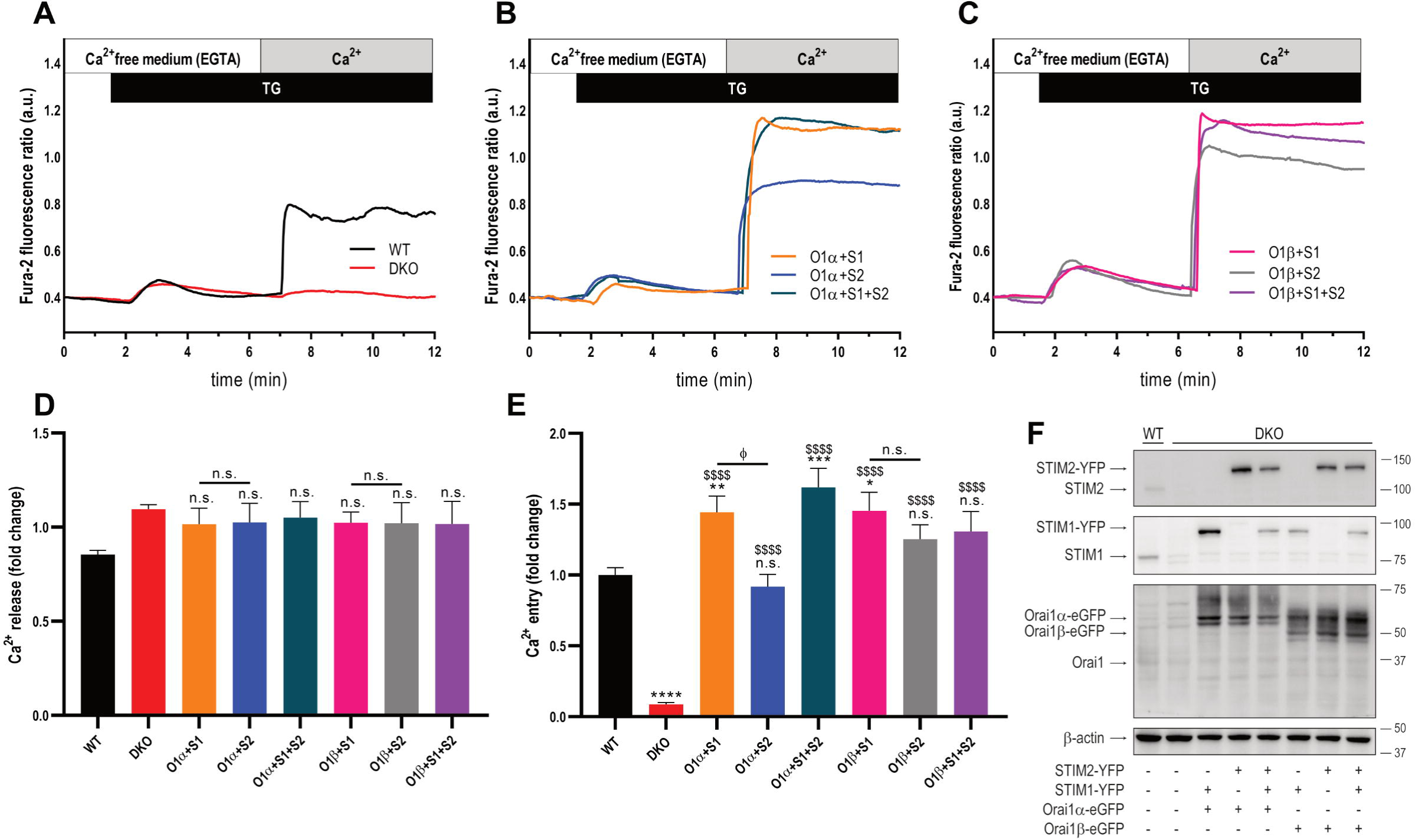
Ca^2+^ entry through the Orai1 variants induced by STIM1 and STIM2. (**A**) Fura-2-loaded wild type HEK-293 cells (WT) and STIM1,2-DKO HEK-293 cells (DKO) were suspended in a Ca^2+^-free (100 µM EGTA) HBS and then stimulated with 2 µM TG followed by reintroduction of external Ca^2+^ (final concentration 1.8 mM) to initiate Ca^2+^ entry. (**B** and **C**) DKO HEK-239 cells were co-transfected with either CMV-driven Orai1α-eGFP (O1α; **B**) or Orai1β-eGFP (O1β; **C**) and either STIM1-YFP (S1), STIM2-YFP (S2) or both (S1+S2) plasmids, as described. Forty-eight hours later cells were loaded with fura-2 and suspended in a Ca^2+^-free HBS (100 µM EGTA added) and then stimulated with TG (2 µM) followed by the addition of extracellular Ca^2+^ (final concentration 1.8 mM) to initiate Ca^2+^ influx. (**D** and **E**) quantification of TG-evoked Ca^2+^ release (**D**) and entry (**E**) determined as described in materials and methods. Bar graphs are represented as mean ± SEM and expressed as fold change over control (WT HEK-293 cells). From left to right, n[=[224, 112, 50, 46, 37, 41, 41 and 23; n values correspond to individual cells. Data were statistically analyzed using Kruskal–Wallis test with multiple comparisons (Dunńs test). * *P* < 0.05, ** *P* < 0.01, \*\*\**P* < 0.001 and \*\*\*\**P* < 0.0001 as compared to WT HEK-293 cells. ^$$^*P* < 0.01 and ^$$$$^*P* < 0.0001 as compared to DKO HEK-293 cells. ^φ^*P* < 0.05 as compared to DKO cells transfected with STIM1 expressing plasmid. (**F**) WT HEK-293 cells (lane 1) and DKO cells either transfected with empty vector or with CMV-driven Orai1α or Orai1β in combination with either STIM1, STIM2 or both plasmids (lanes 2-8) were lysed and then subjected to 10% SDS-PAGE and Western blotting with the anti-STIM2, anti-STIM1 or anti-Orai1 (C-terminal) antibody, as described in material and methods. Membranes were reprobed with the anti-β-actin antibody for protein loading control. Molecular masses indicated on the right were determined using molecular-mass markers run in the same gel. Blots are representative of four separate experiments.

Next, we explored the role of STIM1 and STIM2 on carbachol (CCh)-induced Ca^2+^ mobilization in STIM1/2 DKO HEK-293 cells expressing either Orai1α or Orai1β. In the presence of 1.8 mM extracellular Ca^2+^, CCh (10 µM) elicited Ca^2+^ mobilization. Traces from five representative cells are depicted in Fig. 2A–F. About 99% of WT HEK-293 cells responded to CCh with regenerative Ca^2+^ oscillations (Fig. 2A). In DKO cells, CCh induced a transient rise in the fura-2 fluorescence ratio due to Ca^2+^ release from the intracellular stores and did not respond with Ca^2+^ oscillations (Fig. 2B). The magnitude of Ca^2+^ mobilization induced by CCh was significantly greater in WT, which reveals the role of STIM proteins in CCh-induced Ca^2+^ signals (Fig. 2G; *p* < 0.0001). When DKO cells were co-transfected with CMV-driven Orai1α or Orai1β in combination with STIM1 or STIM2 expression plasmid most cells responded to CCh with a rather sustained increase in the fura-2 fluorescence ratio and Ca^2+^ mobilization was rescued (Fig. 2C-G). We noticed that the Ca^2+^ mobilization induced by CCh in DKO cells expressing Orai1α was greater in the presence of STIM1 than of STIM2 (Fig. 2G; *P*<0.0001). Despite we did not detect any significant difference between STIM1 and STIM2 in cells expressing Orai1β (Fig. 2G), our findings further indicate that STIM1 is more effective than STIM2 mediating CCh-induced Ca^2+^ mobilization.

**Fig. 2.**
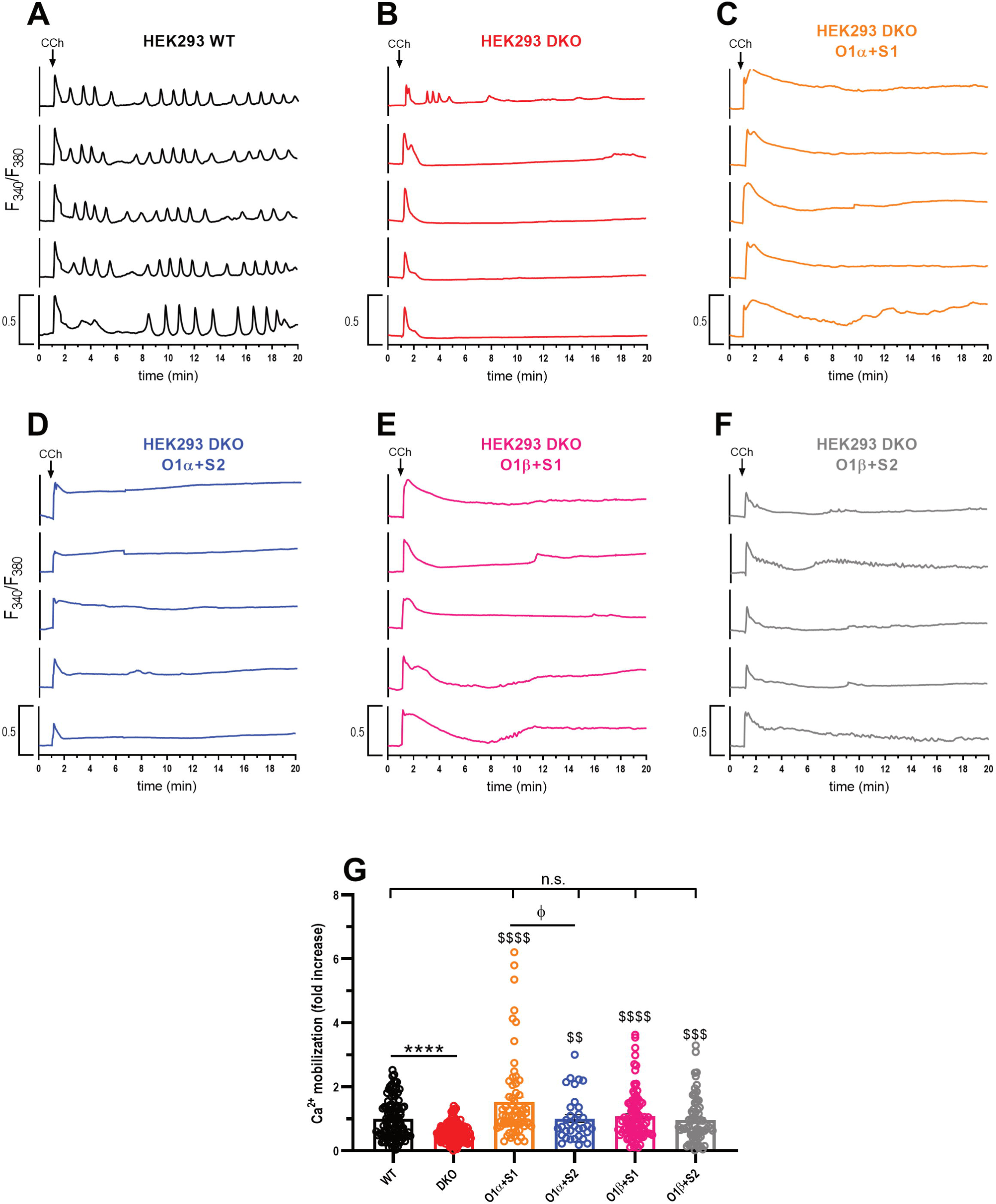
CCh-evoked Ca^2+^ mobilization in cells expressing the Orai1 variants in combination with STIM1 or STIM2. (**A**-**F**) Representative Ca^2+^ mobilization in response to 10[µM CCh measured using fura-2 in wild type HEK-293 cells (WT), STIM1,2-DKO HEK-293 cells (DKO) and DKO cells co-transfected with either CMV-driven Orai1α-eGFP (O1α) or Orai1β-eGFP (O1β) and either STIM1-YFP (S1), STIM2-YFP (S2) or both (S1+S2) plasmids, as described. Cells were superfused with HBS containing 2[mM Ca^2+^ and stimulated with 10[µM CCh at 1[min (indicated by arrow). Representative traces from five cells/condition were chosen to represent the datasets. (**G**) Quantification of Ca^2+^ mobilization for all the conditions from **A** to **F** estimated in all the cells. (for **G**, from left to right, *n*=110, 141, 67, 34, 89 and 70; n-values correspond to individual cells). Scatter plots are represented as mean[±[SEM and expressed as fold change over control (WT HEK-293 cells). Data were statistically analyzed using Kruskal–Wallis test with multiple comparisons (Dunńs test). \*\*\*\**P* < 0.0001 as compared to WT HEK-293 cells. ^$$^*P* < 0.01, ^$$$^*P* < 0.001 and ^$$$$^*P* < 0.0001 as compared to DKO cells. ^φ^*P* < 0.05 as compared to DKO cells transfected with STIM1 expressing plasmid.

We realized that, using expression plasmids with CMV promoter for Orai1 variants and STIM proteins, cells did not respond to CCh with regenerative Ca^2+^ oscillations, mostly due to the overexpression of the target proteins [17]. Hence, we repeated these experimental protocols using plasmids with thymidine kinase (TK) promoter, which yield a protein expression level significantly smaller than plasmids with the CMV promoter [18]. DKO cells were transfected with shOrai1 to reduce the expression of native Orai1 (Fig. 3). Co-transfection with TK-driven Orai1α or Orai1β with STIM1 rescued SOCE in DKO cells (Figs. 3A-E); however, as observed with CMV-driven plasmids, co-transfection of STIM2 with Orai1α was less effective than STIM1 (Figs. 3B, D and E; *P*<0.0001), while the Ca^2+^ signal generated in cells expressing Orai1β was similar in the presence of STIM1 or STIM2 (Fig. 3C, D and F). These findings further indicate that STIM2 is less effective inducing SOCE by Orai1α, but not by Orai1β, as mentioned above using CMV-driven plasmids. Fig. 3F shows the expression of the target proteins under the different experimental conditions. Furthermore, we analyzed the functional role of STIM1 and STIM2 in CCh-evoked Ca^2+^ oscillations using TK-driven plasmids. In the presence of 1.8 mM extracellular Ca^2+^, WT HEK-293 cells responded to CCh with regenerative Ca^2+^ oscillations (Fig. 4A, H, I), with an average of 10.0 ±[0.4 oscillations/20 min (Fig. 4J). In DKO cells, where SOCE is impaired, only 22% of the cells responded with Ca^2+^ oscillations after stimulation with CCh, with an average of 3.2 ± 0.2 oscillations/20 min (Fig. 4B and H). The remaining cells responded with transient Ca^2+^ signals because of Ca^2+^ release from the intracellular stores. As a result, the magnitude of Ca^2+^ mobilization induced by CCh was significantly greater in WT, which further confirms the role of STIM proteins in CCh-induced Ca^2+^ signals (Fig. 4G; *P*<0.0001). When DKO cells were co-transfected with TK-driven Orai1α or Orai1β in combination with STIM1 or STIM2 expression plasmid, the percentage of cells that responded with regenerative Ca^2+^ oscillations enhanced to 72% and 54% (for Orai1α+STIM1 and Orai1α+STIM2, respectively) and 58% and 69% (for Orai1β+STIM1 and Orai1β+STIM2, respectively) (Fig. 4H). Only the combination of the expression of Orai1α+STIM1 in DKO cells resulted in a percentage of oscillating cells that was not statistically different from WT cells (Fig. 4H; *P*<0.0001) but no combination completely rescued the number of oscillations observed in WT cells (Fig. 4J; *P*<0.01). Co-expression of Orai1α and STIM1 in DKO cells completely rescued CCh-induced Ca^2+^ mobilization (Fig 4G), meanwhile co-expression of Orai1β with STIM1 or STIM2 only partially rescued CCh-induced Ca^2+^ mobilization, and expression of Orai1α and STIM2 was unable to rescue Ca^2+^ mobilization induced by CCh in DKO cells as compared to WT cells (Fig. 4G; *P*<0.0001) thus resulting in a Ca^2+^ response than is significantly smaller than that generated by co-expression of Orai1α and STIM1 (Fig. 4G; *P*<0.0001). Altogether, our results indicate that STIM1 is more efficient that STIM2 generating Orai1α-dependent TG-and CCh-evoked Ca^2+^ signals. By contrast, STIM1 and STIM2 exhibit a similar efficiency activating Ca^2+^ signals generated by Orai1β.

**Fig. 3.**
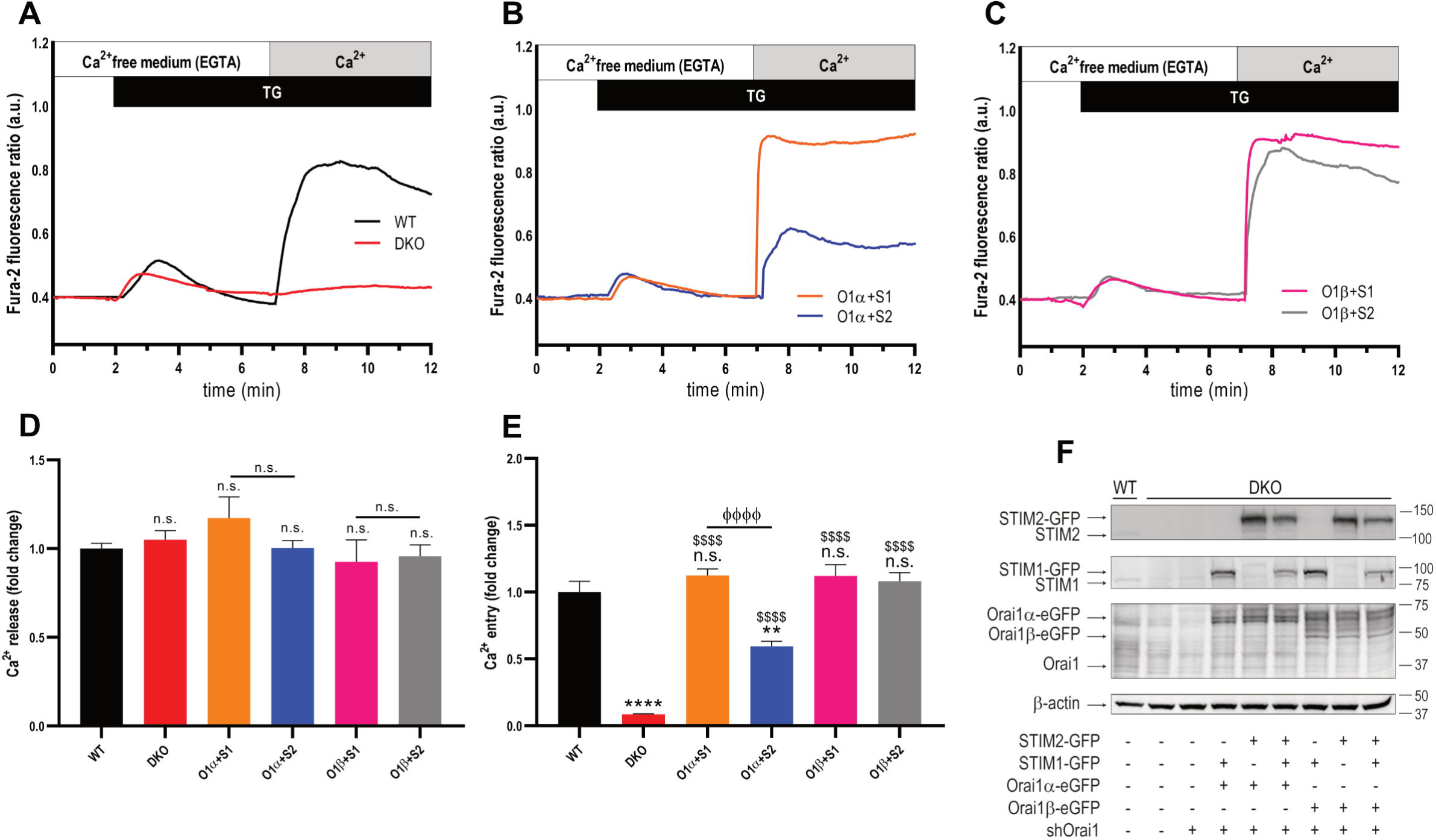
Ca^2+^ entry through the Orai1 variants induced by STIM1 and STIM2 using TK-driven plasmids. (**A**) Fura-2-loaded wild type HEK-293 cells (WT) and STIM1,2-DKO HEK-293 cells (DKO) were suspended in a Ca^2+^-free HBS (100 µM EGTA) and then stimulated with 2 µM TG followed by reintroduction of external Ca^2+^ (final concentration 1.8 mM) to initiate Ca^2+^ entry. (**B** and **C**) DKO cells were co-transfected with either shOrai1 and TK-driven Orai1α-eGFP (O1α; **B**) or Orai1β-eGFP (O1β; **C**) and either STIM1-GFP (S1) or STIM2-GFP (S2) plasmids. Forty-eight hours later cells were loaded with fura-2 and suspended in a Ca^2+^-free HBS (100 µM EGTA added) and then stimulated with TG (2 µM) followed by the addition of extracellular Ca^2+^ (final concentration 1.8 mM) to initiate Ca^2+^ influx. (**D** and **E**) quantification of TG-evoked Ca^2+^ release (**D**) and entry (**E**) determined as described in materials and methods. Bar graphs are represented as mean ± SEM and expressed as fold change over control (WT HEK-293 cells). From left to right, n[=[203, 133, 105, 82, 70 and 109; n values correspond to individual cells. Data were statistically analyzed using Kruskal–Wallis test with multiple comparisons (Dunńs test). \*\**P* < 0.01 and \*\*\*\**P* < 0.0001 as compared to WT HEK-293 cells. ^$$$^*P* < 0.001 and ^$$$$^*P* < 0.0001 as compared to DKO cells. ^φφφ^*P* < 0.001 and ^φφφφ^*P* < 0.0001 as compared to DKO cells transfected with STIM1 expressing plasmid. (**F**) WT HEK-293 cells (lane 1) and DKO cells either transfected with empty vector or with TK-driven Orai1α or Orai1β in combination with either STIM1, STIM2 or both plasmids (lanes 2-8), as described, were lysed and then subjected to 10% SDS-PAGE and Western blotting with the anti-STIM2, anti-STIM1 or anti-Orai1 (C-terminal) antibody, as described in material and methods. Membranes were reprobed with the anti-β-actin antibody for protein loading control. Molecular masses indicated on the right were determined using molecular-mass markers run in the same gel. Blots are representative of four separate experiments.

**Fig. 4.**
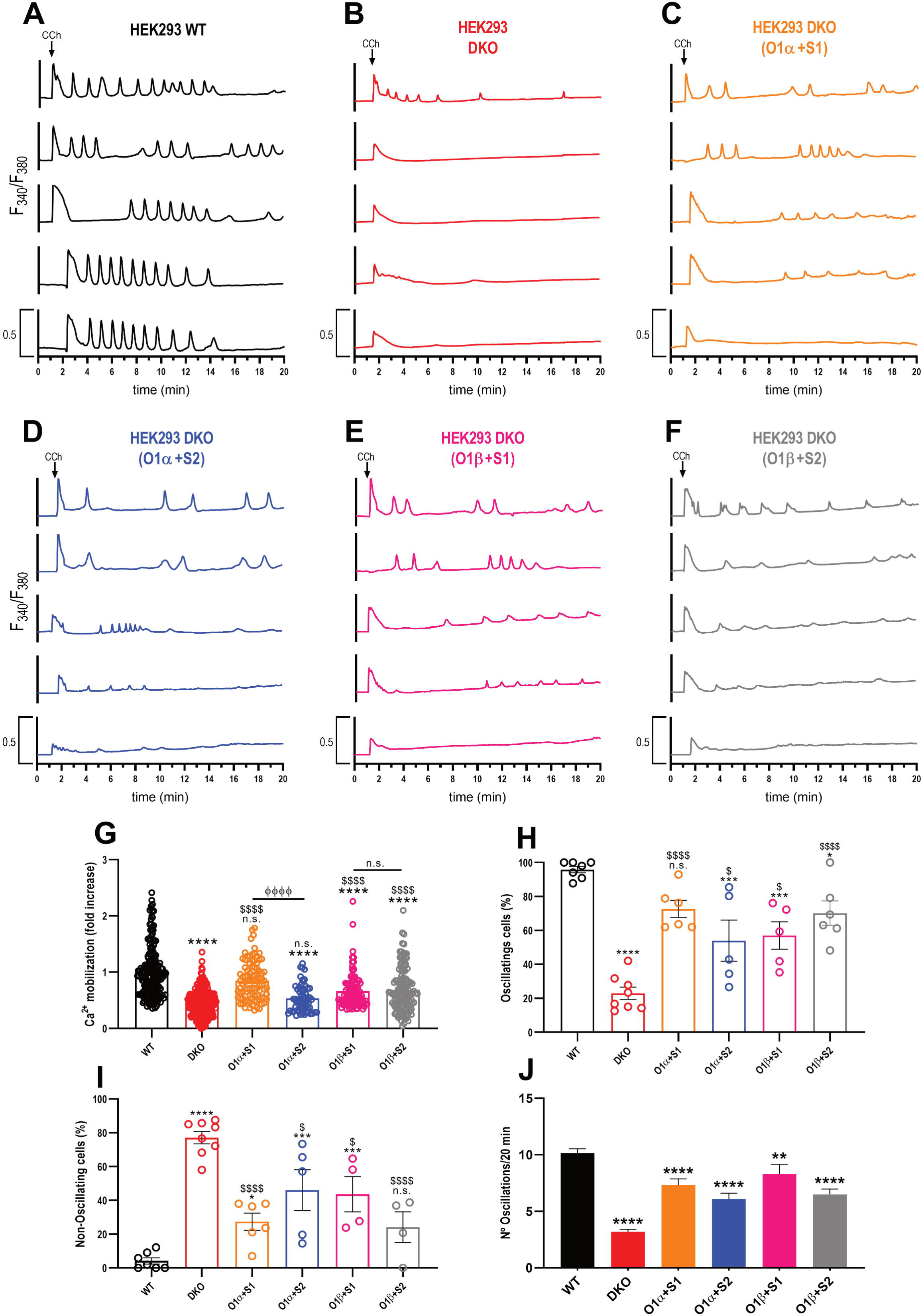
CCh-evoked Ca^2+^ oscillations in cells expressing the Orai1 variants in combination with STIM1 or STIM2. (**A-F**) Representative Ca^2+^ mobilization in response to 10[µM CCh measured using fura-2 in wild type HEK-293 cells (WT), STIM1,2-DKO HEK-293 cells (DKO) and DKO cells co-transfected with either shOrai1 and TK-driven Orai1α-eGFP (O1α) or Orai1β-eGFP (O1β) and either STIM1-GFP (S1) or STIM2-GFP (S2) plasmids, as described. Cells were superfused with HBS containing 2[mM Ca^2+^ and stimulated with 10[µM CCh at 1[min (indicated by arrow). Representative traces from five cells/condition were chosen to represent the datasets. (**G**) Quantification of Ca^2+^ mobilization for all the conditions from **A** to **F** estimated in all the cells. (for **G**, from left to right, *n*=187, 286, 91, 61, 110 and 126; n-values correspond to individual cells). (**H-J**) Quantification of the percentage of oscillating cells (**H**), non-oscillating cells (**I**) and total oscillations/cell in 20 min (**J**) for data presented in **A**–**F** (for **H** to **J**, from left to right, n[=[8, 8, 6, 5, 5 and 6; n values correspond to independent experiments). Scatter plots are represented as mean[±[SEM and expressed as fold change over control (WT HEK-293 cells). Data were statistically analyzed using Kruskal–Wallis test with multiple comparisons (Dunńs test). \**P* < 0.05, \*\**P* < 0.01, \*\*\**P* < 0.001 and \*\*\*\**P* < 0.0001 as compared to WT HEK-293 cells. ^$^*P* < 0.05, and ^$$$$^*P* < 0.0001 as compared to DKO cells. ^φφφφ^*P* < 0.0001 as compared to DKO cells transfected with STIM1 expressing plasmid.

### Interaction of Orai1 variants with STIM1 and STIM2

STIM proteins associate with and activate Orai channels to mediate SOCE. Here we have used confocal Förster Resonance Energy Transfer (FRET) to examine the interaction of STIM1 and STIM2 with Orai1 variants in living HEK293 cells. Co-expression of CMV-driven C-terminally tagged Orai1α-eGFP (Fig. 5A and C) or C-terminally tagged Orai1β-eGFP (Fig. 5B and D) together with CMV-driven C-terminally tagged STIM1-mCherry (Fig. 5A and B) or STIM2-mCherry (Fig. 5C and D), revealed substantial PM (Orai1α-eGFP and Orai1β-eGFP) or intracellular localization (STIM1-mCherry and STIM2-mCherry). A mean nFRET value of 0.106 ± 0.002, 0.095 ± 0.003, 0.159 ± 0.009 and 0.074 ± 0.001 suggests a direct coupling of STIM1-Orai1α, STIM1-Orai1β, STIM2-Orai1α and STIM2-Orai1β, respectively, while the cell is in a quiescent state. One of the functions attributed to STIM2 is its ability to activate Orai1 in order to counteract minor leakage from the ER [29]. It is noteworthy that, in a quiescent cell, the interaction between STIM2 and Orai1α is significantly more pronounced compared to the other studied conditions (Fig. 5A-D, control, and 5E control; *P*<0.0001). As expected, stimulation with 2 µM TG for 1 min in the presence of 1.8 mM extracellular Ca^2+^ resulted in a significant increase in the mean nFRET value in all tested combinations, STIM1-Orai1α: 0.378 ± 0.030; STIM1-Orai1β: 0.334 ± 0.037; STIM2-Orai1α: 0.269 ± 0.018; and STIM2-Orai1β: 0.114 ± 0.002. Our data show a strong association between STIM1 and both Orai1 variants or STIM2 and Orai1α (Fig. 5A-C, TG, and 5E, TG; *P*<0.0001), while a weaker association between STIM2 and Orai1β upon TG stimulation was observed (Fig. 5D, TG, and 5E, TG; *P*<0.05).

**Fig. 5.**
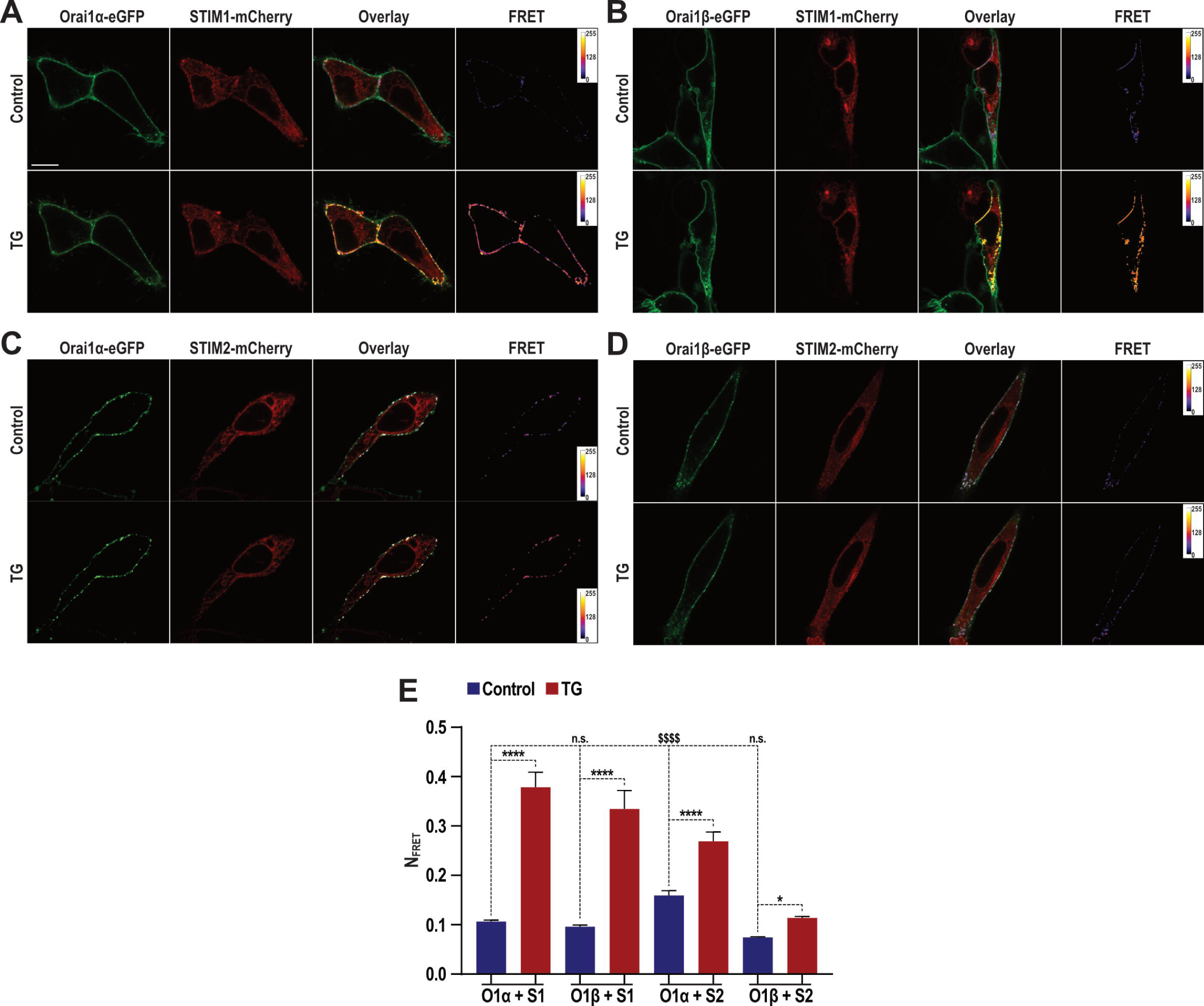
*In vivo* interaction of STIM1 and STIM2 with Orai1 variants. (**A-D**) Resting (control) and stimulated (2 µM TG for 1 min) WT HEK293 cells co-expressing Orai1α-eGFP (**A** and **C**) or Orai1β-eGFP (**B** and **D**) with STIM1-mCherry (**A** and **B**) or STIM2-mCherry (**C** and **D**), their overlay (**A-D**, Overlay) and calculated FRET values (**A-D**, FRET) are presented. Cells were superfused with a medium containing 1.8 mM Ca^2+^. The FRET values depicted in **E** were calculated from the averages of whole cell areas determined from the respective number of cells showing significantly increased FRET values. Values are mean ± S.E. of twenty independent experiments. Data were statistically analyzed using Kruskal–Wallis test with multiple comparisons (Dunńs test). \**P* < 0.05, \*\*\*\**P* < 0.0001, ^$$$$^*P* < 0.0001 as compared to their respective controls (untreated cells). The scale bar in the box represents 10 µm.

We have further investigated the interaction of Orai1 variants with STIM1 and STIM2 by looking for coimmunoprecipitation between the target proteins from cell lysates. Immunoprecipitation and subsequent SDS-PAGE and Western blotting were conducted using DKO HEK-293 cells co-transfected with CMV-driven Orai1α or Orai1β in combination with either STIM1, STIM2 or both STIM proteins. The experiments were performed both at resting conditions and after stimulation with 2 μM TG for 1 min in the presence of 1.8 mM extracellular Ca^2+^. As revealed by the FRET experiments, TG stimulated a significant coimmunoprecipitation between Orai1α and Orai1β with STIM1 and STIM2 (Fig. 6A-D, *P*<0.05). Western blotting with anti-Orai1 antibody confirmed a similar content of this protein in all lanes (Fig. 6A and B). Next, we explored the interaction of the Orai1 variants with STIM1 and STIM2 when both STIM proteins were expressed. As shown in Fig. 6, the interaction of STIM1 with Orai1α or Orai1β was similar when transfected alone or in combination with STIM2, and similar results were obtained when we analyzed the association of STIM2 with the Orai1 variants in the absence or presence of STIM1.

**Fig. 6.**
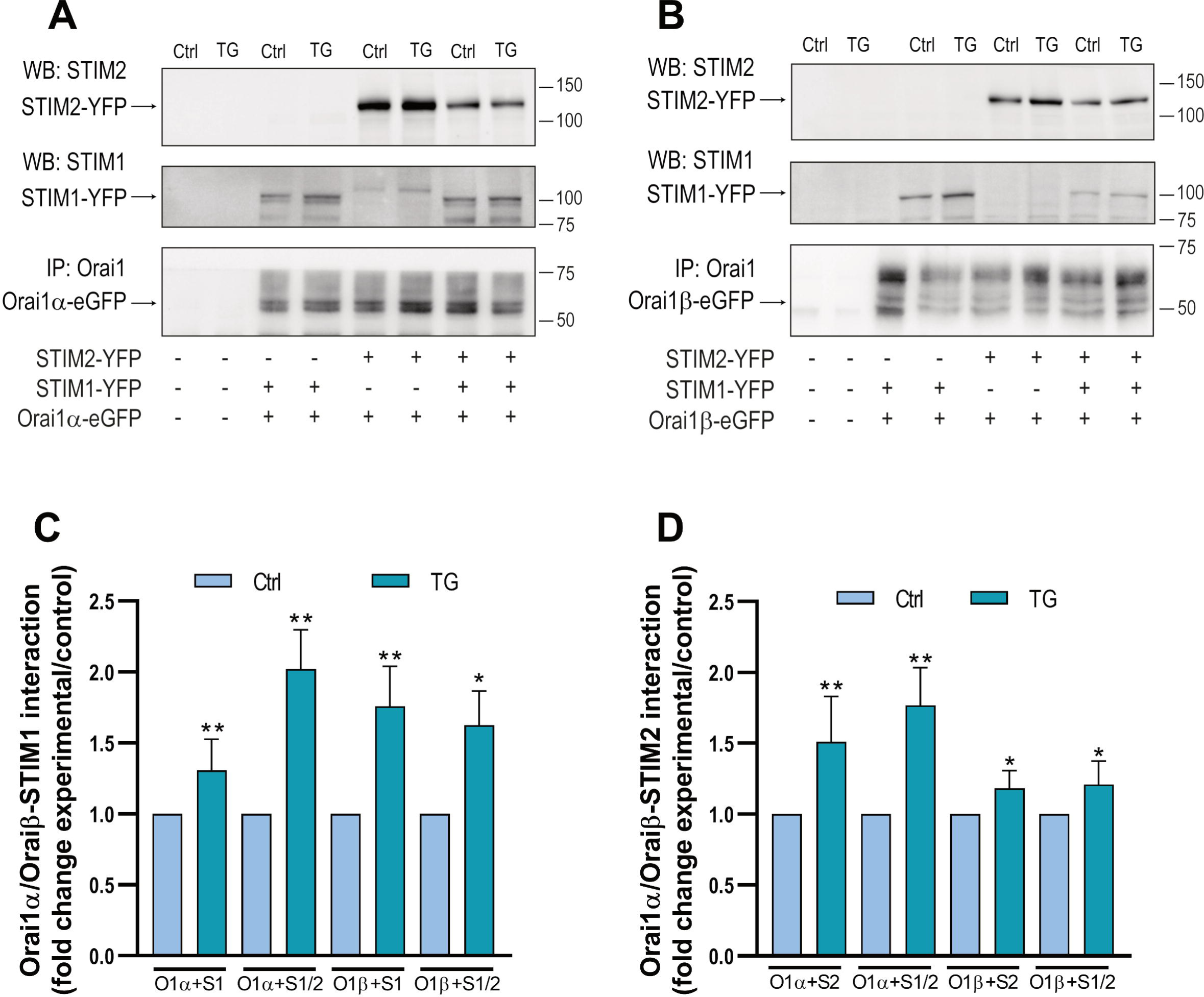
Interaction of Orai1α and Orai1β with STIM1 and STIM2. STIM1,2-DKO HEK-293 cells were left untreated (**A**, lanes 1 and 2) or co-transfected with CMV-driven Orai1α-eGFP (**A**, lanes 3-8) or Orai1β-eGFP (**B**, lanes 3-8) in combination with either STIM1-YFP, STIM2-YFP or both plasmids. Forty-eight hours later cells were then treated in the absence or presence of 2 µM TG for 1 min and lysed. Cell lysates were then subjected to 10% SDS-PAGE and Western blotting with the anti-STIM2, anti-STIM1 or anti-Orai1 (C-terminal) antibody, as described in material and methods. Membranes were reprobed with the anti-β-actin antibody for protein loading control. Molecular masses indicated on the right were determined using molecular-mass markers run in the same gel. Blots are representative of four to five separate experiments. (**C-D**) Quantification of the interaction of STIM1 (**C**) or STIM2 (**D**) interaction with Orai1α or Orai1β. Data were statistically analyzed using Mann–Whitney U test. \**P* < 0.05 and \*\**P* < 0.01 as compared to their respective controls (untreated cells).

### Plasma membrane expression of Orai1α and Orai1β in the absence and presence of STIM1 and/or STIM2

We and others have previously demonstrated that a pool of compartmentalized Orai1 translocates to the PM upon Ca^2+^ store depletion [19, 21]. We have analyzed the PM expression of Orai1 variants by biotinylation of surface proteins. Experiments were performed in STIM1/2 DKO HEK-293 cells co-transfected with CMV-driven Orai1α or Orai1β in combination with either STIM1, STIM2 or both STIM proteins. The experiments were performed both at resting conditions and after stimulation with 2 μM TG for 1 min in the presence of 1.8 mM extracellular Ca^2+^. As shown in Fig. 7, PM expression of Orai1α and Orai1β was detected in resting cells, and stimulation with 2[µM TG significantly enhanced the amount of Orai1α resident in the PM (*P*<0.0001; n[=[6). Interestingly, expression of STIM1, STIM2 or both significantly attenuated the PM expression of Orai1α and Orai1β, both at resting conditions and after stimulation with TG, despite the surface expression of both Orai1 variants upon treatment with TG was significantly greater than in resting cells (Fig. 7; *P*<0.05). Altogether, these findings indicate that expression of STIM1 and/or STIM2 significantly attenuates the surface expression of Orai1 variants.

**Fig. 7.**
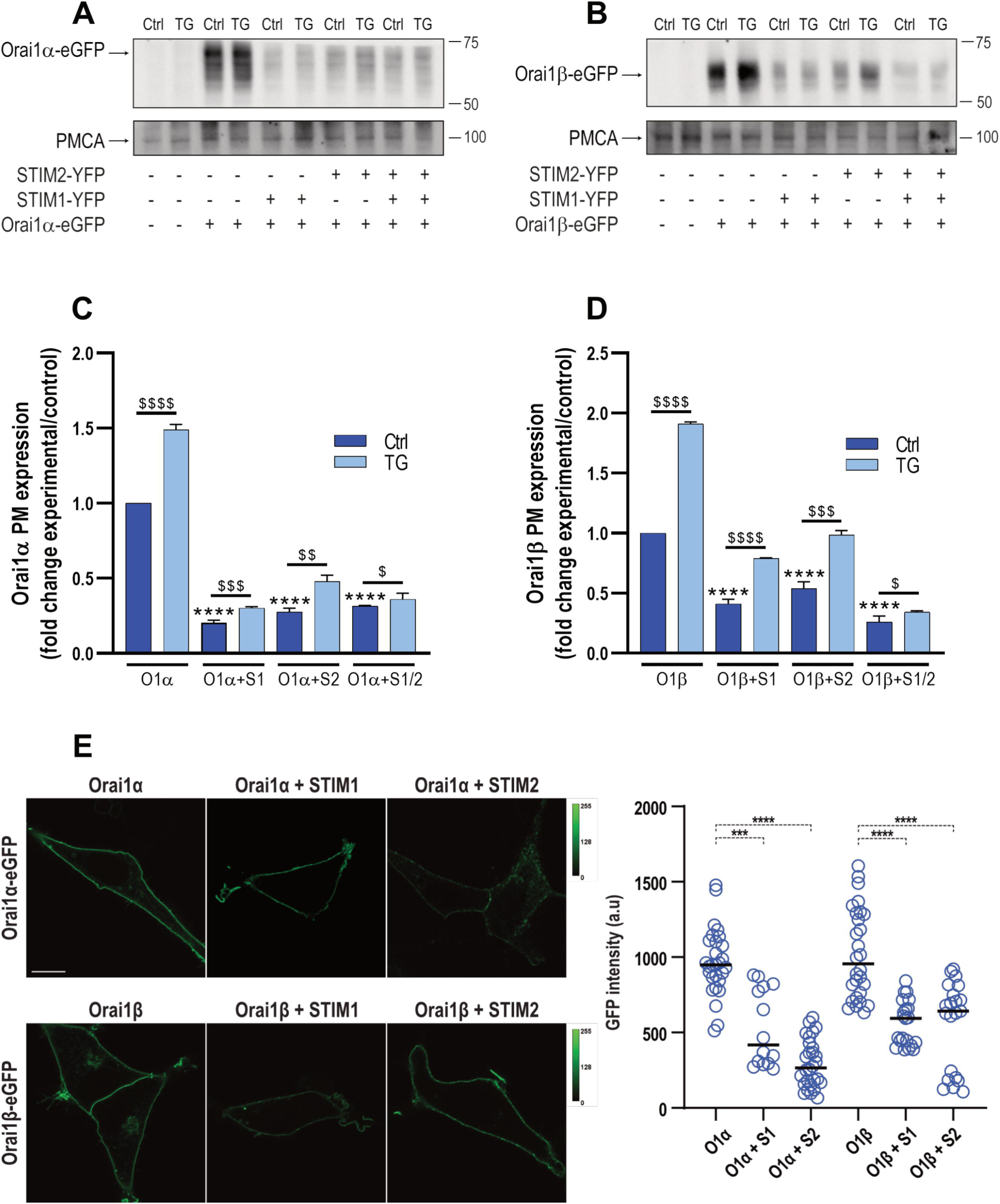
Plasma membrane expression of Orai1α and Orai1β upon co-expression with STIM1 and/or STIM2. STIM1,2-DKO HEK-293 cells were left untreated (panels **A** and **B**, lanes 1 and 2) or co-transfected with CMV-driven Orai1α-eGFP (**A**, lanes 3-10) or Orai1β-eGFP (**B**, lanes 3-10) alone or in combination with either STIM1-YFP, STIM2-YFP or both plasmids, as indicated. Forty-eight hours later cells were then treated in the absence (Ctrl) or presence of 2 µM TG for 1 min and lysed. Samples were then mixed with biotinylation buffer containing EZ-Link sulfo-NHS-LC-biotin, and cell surface proteins were labeled by biotinylation. Labeled proteins were pulled down with streptavidin-coated agarose beads. The pellet (containing the plasma membrane fraction) and the supernatant were analyzed by SDS-PAGE and Western blot analysis using anti-Orai1 C-terminal antibody, as indicated. Membranes were probed with anti-PMCA antibody. Molecular masses indicated on the right were determined using molecular-mass markers run in the same gel. These results are representative of 6 separate experiments. (**C-D**) Quantification of Orai1 plasma membrane expression under the different experimental conditions normalized to the PMCA expression is depicted in the bar graph. Data are represented as mean[±[SEM and expressed as fold increase (experimental/control). Data were statistically analyzed using Kruskal–Wallis test with multiple comparisons (Dunn’s test). ^$^*P* < 0.05, ^$$^*P* < 0.01, ^$$$^*P* < 0.001 and ^$$$$^*P* < 0.0001 as compared to their respective control (untreated cells). \*\*\*\**P* < 0.0001 as compared to the Orai1 expression in the absence of STIM proteins. (**E**) HEK-293 cells were transfected with CMV-driven Orai1α-eGFP or Orai1β-eGFP alone or in combination with CMV-driven STIM1-mCherry or STIM2-mCherry. Forty-eight hours later GFP fluorescence was detected using an LSM900 confocal microscope. The images show representative confocal images of Orai1α-eGFP or Orai1β-eGFP. The scale bar represents 10 μm. Scatter plots represent GFP intensity at the different experimental conditions. Data are presented as mean ± SEM and are statistically analyzed using Kruskal–Wallis test with multiple comparisons (Dunńs test). \*\*\**P* < 0.001 and \*\*\*\**P* < 0.0001 as compared to HEK-293 cells transfected with Orai1α-eGFP or Orai1β-eGFP alone.

The PM expression of Orai1 variants at basal conditions was further analyzed in HEK-293 cells transfected with CMV-driven Orai1α-eGFP or Orai1β-eGFP alone or in combination with CMV-driven STIM1 or STIM2 by analyzing the GFP fluorescence at the PM. GFP fluorescence measurements were carried out on an LSM900 confocal microscope. As shown in Fig. 7E, upon expression of Orai1α or Orai1β GFP fluorescence was detected evenly distributed at the PM. Measurement of GFP fluorescence at the PM revealed that the GFP intensity was significantly smaller upon the expression of STIM1 and STIM2 than when Orai1α or Orai1β were expressed independently. These findings confirm the results obtained by biotinylation and further indicate that the expression of STIM1 and STIM2 significantly attenuates the PM expression of Orai1α or Orai1β.

### STIM1 and STIM2 regulates the expression of Orai1α and Orai1β

The results presented in Fig. 7 indicate that the expression of Orai1α and Orai1β at the PM is significantly reduced upon co-expression of STIM1 or STIM2. Hence, we have further analyzed whether STIM1 and STIM2 regulate the cellular expression of Orai1α and Orai1β at the protein level. To investigate this issue, STIM1/2 DKO HEK-293 cells were co-transfected with Orai1α or Orai1β in combination with STIM1 or STIM2 using CMV-driven plasmids with Orai1:STIM ratios 1:1 and 1:2, and we compared levels of expression of Orai1 variants by Western blotting (Fig. 8A and B). Fig. 8C depicts the expression of STIM1 and STIM2 when cells were transfected with 1 µg/mL STIM plasmid (for 1:1 Orai1:STIM ratio) or 2 µg/mL STIM plasmid (for 1:2 Orai1:STIM ratio). As shown in Fig. 8A, DKO cells transfection with Orai1α in combination with STIM1 led to a significant drop in Orai1α expression both at the Orai1:STIM1 ratio of 1:1 and 1:2, being significantly smaller at the 1:2 ratio (*P*<0.05; *n*=6). Similar results were obtained when cells were co-transfected with Orai1α and STIM2. Furthermore, expression of STIM1 or STIM2 at 1:1 and 1:2 expression ratios significantly reduced the extent of Orai1β expression, with a significantly greater effect when STIM proteins were co-transfected at an Orai1:STIM ratio of 1:2 (Fig. 8B; *P*<0.05; *n*=6). These findings, indicate that Orai1 variant expression is modulated by the protein level of STIM1 and STIM2.

**Fig. 8.**
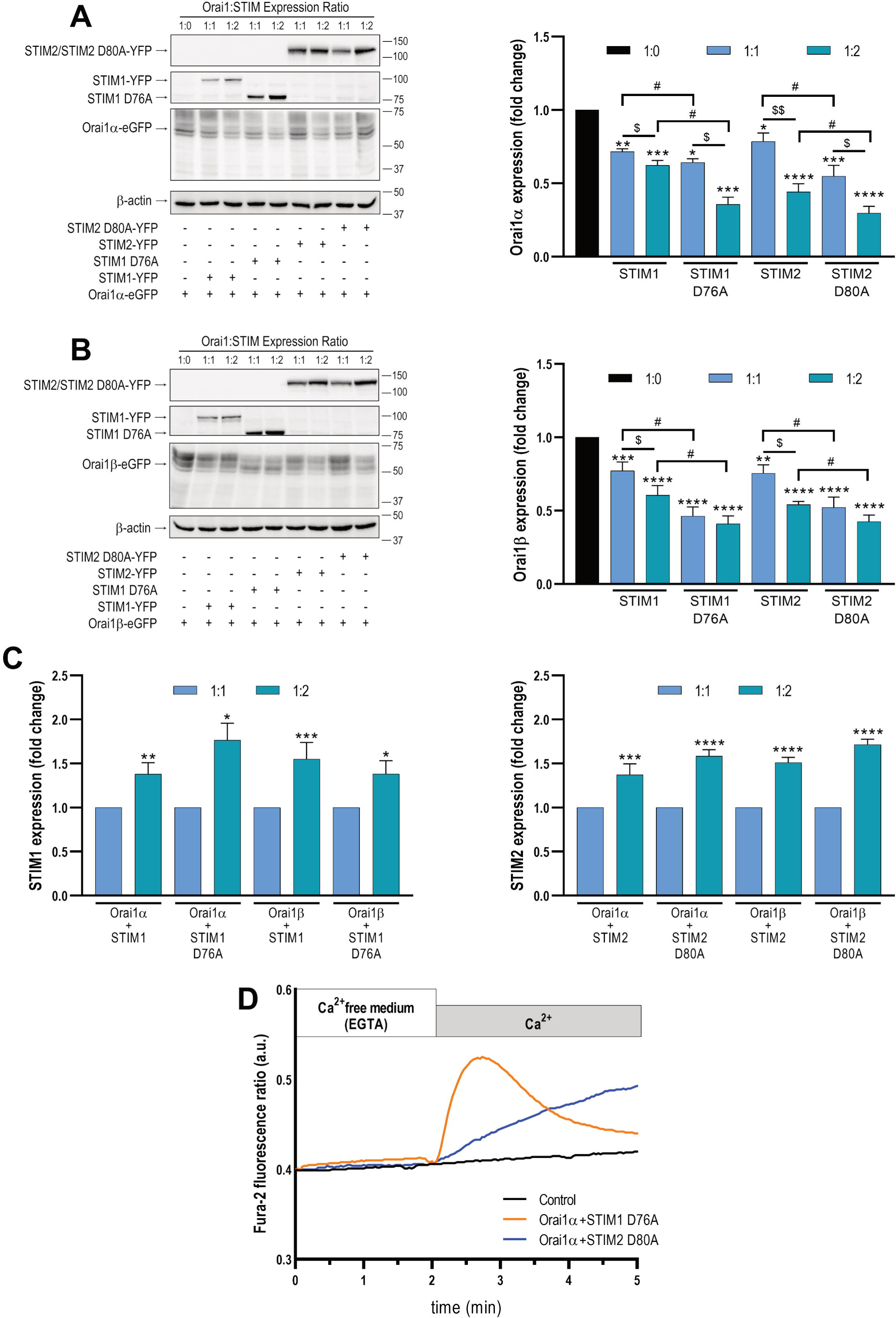
STIM1 and STIM2 modulate the expression of Orai1α and Orai1β. STIM1,2-DKO HEK-293 cells were co-transfected with CMV-driven Orai1α (**A**) or Orai1β (**B**) in combination with either STIM1-YFP, STIM2-YFP or the STIM1-D76A and STIM2-D80A-YFP mutants, with a relative Orai1:STIM expression ratio of 1:1 or 1:2, as indicated. Forty-eight hours later cells were lysed, and cell lysates were analyzed by SDS-PAGE and Western blot analysis using anti-Orai1 C-terminal antibody. Membranes were probed with anti-STIM1, anti-STIM2 or anti β-actin antibody (**A-C**). Molecular masses indicated on the right were determined using molecular-mass markers run in the same gel. These results are representative of 6 separate experiments. Bar graphs represent the quantification of Orai1α, Orai1β, STIM1 and STIM2 expression under the different experimental conditions normalized to the β-actin expression. Data are represented as mean[±[SEM and expressed as fold increase (experimental/control). (**A-B**) Data were statistically analyzed using Kruskal–Wallis test with multiple comparisons (Dunn’s test). \**P* < 0.05, \*\**P* < 0.01, \*\*\**P* < 0.001 and \*\*\*\**P* < 0.0001 as compared to the expression of Orai1α or Orai1β in the absence of STIM. ^$^*P* < 0.05 and ^$$^*P* < 0.01 as compared to the Orai1 expression at Orai1:STIM expression ratio of 1:1. ^#^*P* < 0.05 as compared to the Orai1 expression in the presence of WT STIM proteins. (**C**) Data were statistically analyzed using Mann-Whitney U test. \**P* < 0.05, \*\**P* < 0.01, \*\*\**P* < 0.001 and \*\*\*\**P* < 0.0001 as compared to the expression of STIM1 or STIM2 at Orai1:STIM relative expression ratio of 1:1. (**D**). STIM1,2-DKO HEK-293 cells were co-transfected with CMV-driven Orai1α in combination with either STIM1 D76A or STIM2-D80A mutant, with empty vector (Control). Forty-eight hours later cells were loaded with fura-2 and suspended in a Ca^2+^-free HBS (100 µM EGTA added). Ca^2+^ was added to the extracellular medium at a final concentration of 1.8 mM to initiate Ca^2+^ influx.

As shown above in Fig. 5, there is some STIM-Orai1 interaction at resting conditions. To further explore whether the interaction of Orai1 with STIM proteins modulates the Orai1α and Orai1β expression, we co-transfected cells with Orai1α or Orai1β in combination with the D76A EF-hand mutant of STIM1 and the equivalent D80A mutant of STIM2, which have been reported to disrupt Ca^2+^ binding and result in constitutive STIM1 or STIM2 activation, respectively [30, 31] (Fig. 8D). As depicted in Fig. 8A and B, the protein level of Orai1α and Orai1β was significantly reduced when co-expressed with the mutant forms of STIM1 and STIM2 as compared to co-expression with WT STIM proteins, either at 1:1 or 1:2 expression ratios. These findings strongly suggest that the interaction with STIM1 and STIM2 modulate Orai1α and Orai1β protein content.

Using the same CMV-promoter Orai1α-EGFP and Orai1β-EGFP expression plasmids, and under the same experimental conditions, in a previous study we did not detect any interference in the expression of Orai1α or Orai1β when co-expressed together or in combination with other intracellular proteins such as ARF6 [21], which suggests that, with the experimental procedure used, the protein transcription and translation is not significantly affected by co-expression with other plasmids. To further confirm that the expression of Orai1α and Orai1β is regulated by STIM expression we analyzed the level of expression of native Orai1α and Orai1β after silencing or overexpression of STIM1 and STIM2. As depicted in Fig. 9A, detection of STIM1, STIM2 and Orai1 in WT HEK-293 by Western blotting revealed a single protein band for STIM2 and STIM1 at the predicted molecular weights of 85 and 75 kDa, respectively, and several diffuse bands for Orai1 around 37 kDa, as previously described [11]. The multiple bands corresponding to Orai1 disappeared after treatment with PNGase F, which hydrolyzes glycosylamine linkage of asparagine-linked oligosaccharides, leading two bands corresponding to Orai1α and Orai1β (Fig. 9A). Our results indicate that overexpression of STIM1 or STIM2 significantly reduced the protein level of Orai1α and Orai1β (Fig. 9B-E; *P*<0.01; n[=[6). By contrast, attenuation of STIM1 and STIM2 expression significantly enhanced the protein content of native Orai1α and Orai1β (Fig. 9B-E; *P*<0.05; n[=[6). These findings provide further evidence for the regulation of Orai1α and Orai1β protein content by STIM1 and STIM2.

**Fig. 9.**
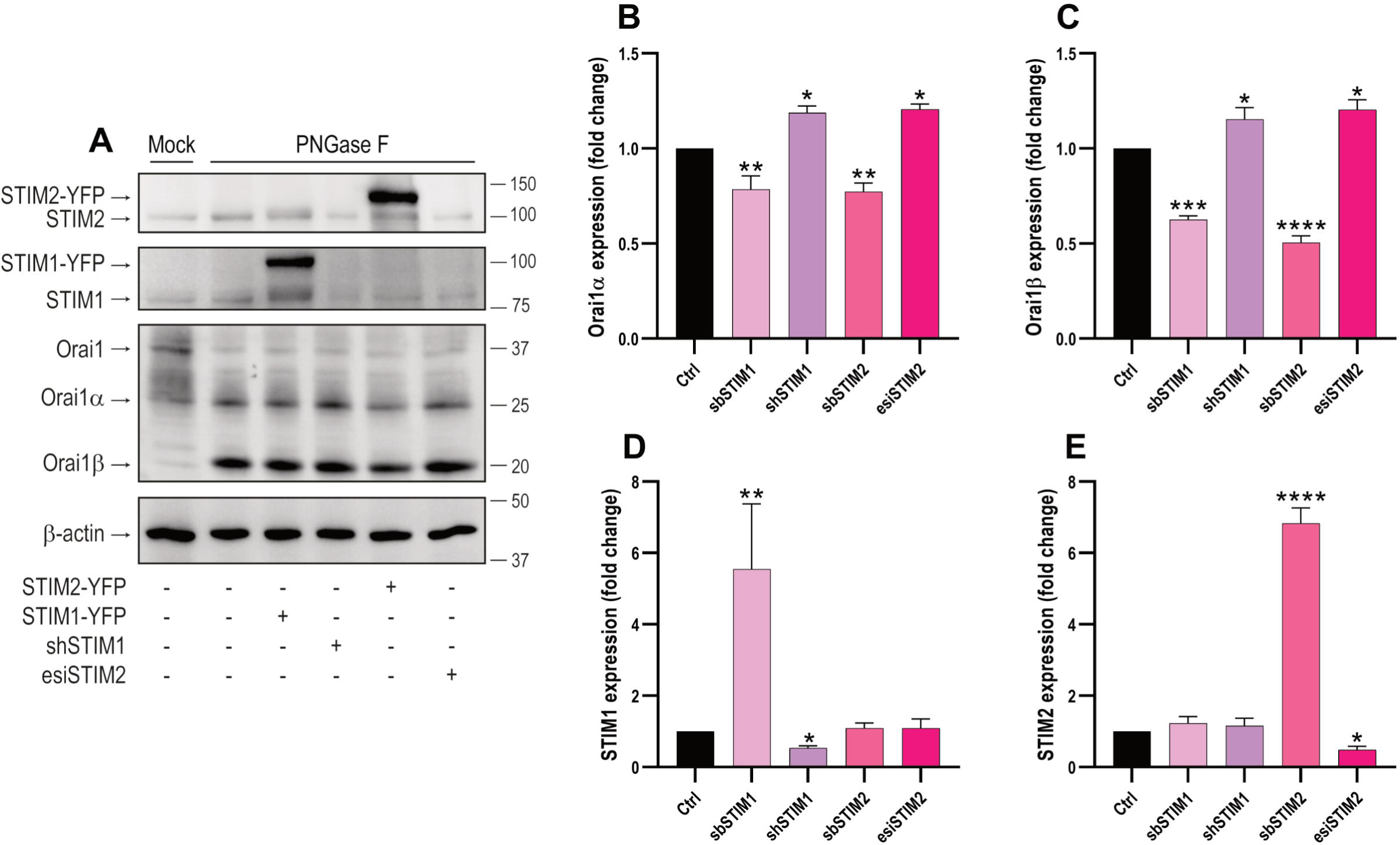
Effect of STIM1 and STIM2 expression on the Orai1α and Orai1β protein level. (A) HEK-293 cells were transfected with empty vector (Ctrl), STIM1-YFP or STIM2-YFP expression plasmids or with shSTIM1 or esiSTIM2, as indicated. Forty-eight hours later cells lysed. Cell lysates were treated with PNGaseF and then subjected to 10% SDS-PAGE and Western blot analysis using anti-STIM1, anti-STIM2 or anti-Orai1 C-terminal antibody. Membranes were reprobed with anti β-actin antibody for protein loading control. Molecular masses indicated on the right were determined using molecular-mass markers run in the same gel. These results are representative of 6 separate experiments. (**B-E**) Quantification of Orai1α, Orai1β, STIM1 and STIM2 expression under the different experimental conditions normalized to the β-actin expression. Data are represented as mean[±[SEM and expressed as fold change (experimental/control). Data were statistically analyzed using Kruskal–Wallis test with multiple comparisons (Dunn’s test). \**P* < 0.05, \*\**P* < 0.01, \*\*\**P* < 0.001 and \*\*\*\**P* < 0.0001 as compared to the protein expression in WT cells.

The protein expression is regulated both by the synthesis rate and the stability of the translated protein. In order to discern between these possibilities, we explored the effect of STIM expression on Orai1α and Orai1β turnover in the presence of the protein synthesis inhibitor cycloheximide (CHX). Cells were transfected with Orai1α or Orai1β alone or in combination with STIM1 and STIM2 and treated with CHX or the vehicle. Protein expression was determined seven hours later. As depicted in Fig. 10A, Orai1α protein expression fell within seven hours by 70, 56 and 60% after co-expression with STIM1, STIM2 or both, respectively (*P*<0.001; n[=[6). CHX by itself attenuated the expression of Orai1α and Orai1β in the absence of STIM proteins by 30±5 and 29±8%, respectively. In the presence of CHX the expression of Orai1α decreased after co-expression with STIM1, STIM2 or both by 78, 75 and 69%, respectively, relative to the expression of Orai1α alone in the presence of CHX (Fig. 10A; *P*<0.0001; n[=[6). Similar results were obtained with Orai1β (Fig. 10B; *P*<0.0001; n[=[6). The drop in Orai1 variant expression was similar in the absence or presence of CHX which indicates that protein synthesis is not involved in the regulation of Orai1 expression by STIM proteins. In mammalian cells two major pathways for protein degradation includes the ubiquitin-proteasome pathway and the lysosomal-dependent proteolysis. To further ascertain whether the drop in Orai1 expression by STIM is mediated by protein degradation and discern among the degradation pathways we treated cells with MG132, a proteasome inhibitor [32], and bafilomycin A1, an inhibitor of the vacuolar H^+^-ATPase that prevents lysosomal protein degradation [33]. Degradation of the Orai1 variants was similar in the absence and presence of MG132 (Fig. 10C and D; *n*=6), which indicates that proteasomal degradation is not involved in the modulation of Orai1 expression by STIM proteins. Interestingly, Orai1α and Orai1β protein loss in the presence of STIM1 and STIM2 was significantly reduced by treatment with bafilomycin A1 (Fig. 10E and F; *n*=6), which demonstrates that STIM proteins modulate Orai1 variant expression level by endo-lysosomal degradation.

**Fig. 10.**
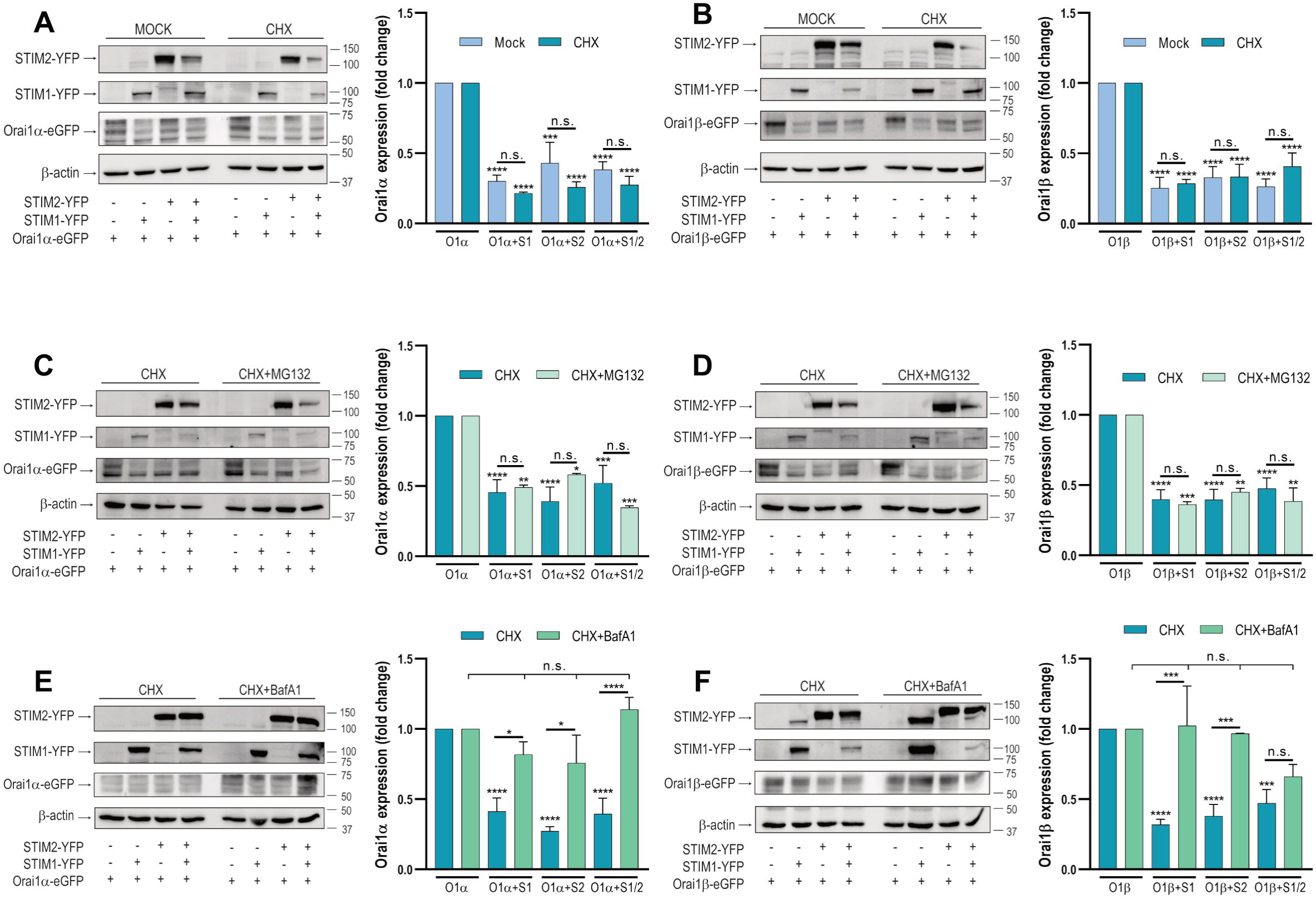
STIM1 and STIM2 regulate lysosomal degradation of Orai1α and Orai1β. STIM1,2-DKO HEK-293 cells were co-transfected with CMV-driven Orai1α-eGFP (**A, C, E**) or Orai1β-eGFP (**B, D, F**) in combination with either STIM1-YFP, STIM2-YFP or both plasmids, as indicated. Forty-eight hours later cells were either treated with cycloheximide (CHX; 10 µM) or the vehicle for 7 h and lysed (**A**-**B**) or were treated with cycloheximide (10 µM) alone or in combination with 10 µM MG132 for 7 h and lysed (**C**-**D**) or were treated with cycloheximide (10 µM) alone or in combination with 1 µM bafilomycin A1 (BafA1) for 7 h and lysed (**E**-**F**). Cell lysates were analyzed by SDS-PAGE and Western blot analysis using anti-STIM1, anti-STIM2 or anti-Orai1 C-terminal antibody. Membranes were reprobed with anti β-actin antibody. Molecular masses indicated on the right were determined using molecular-mass markers run in the same gel. These results are representative of 6 separate experiments. Bar graphs show quantification of Orai1α and Orai1β expression under the different experimental conditions normalized to the β-actin expression. Data are represented as mean[±[SEM and expressed as fold change (experimental/control). Data were statistically analyzed using Kruskal–Wallis test with multiple comparisons (Dunn’s test). \**P* < 0.05, \*\**P* < 0.01, \*\*\**P* < 0.001 and \*\*\*\**P* < 0.0001 as compared to the Orai expression in the absence of STIM proteins.

### Degradation of Orai1α and Orai1β by STIMs is independent on Ca^2+^ influx through Orai1 but requires protein-protein interaction

We have further explored whether Orai1 degradation requires physical interaction with STIM and the subsequent Ca^2+^ influx through the channel. To address whether Ca^2+^ influx through the channel is involved in the Orai1 degradation we have analyzed the role of STIM1 and STIM2 in the protein content of the pore-dead Orai1-E106Q mutant (Orai1dn), which acts as a dominant-negative channel [34]. Fig. 11A compares the protein content of Orai1α and the Orai1dn mutant when expressed alone or in combination with STIM1 and STIM2. Our results indicate that the expression of STIM1 and STIM2 significantly attenuated the protein level of Orai1α and the Orai1dn mutant (*P*<0.001), with no significant differences detected between the WT and mutant forms of Orai1α (Fig. 11A). Fig. 11B depicts lack of TG-induced Ca^2+^ influx in cells expressing Orai1dn. These findings strongly indicates that Ca^2+^ influx through the channel is not essential for its degradation evoked by STIM proteins.

**Fig. 11.**
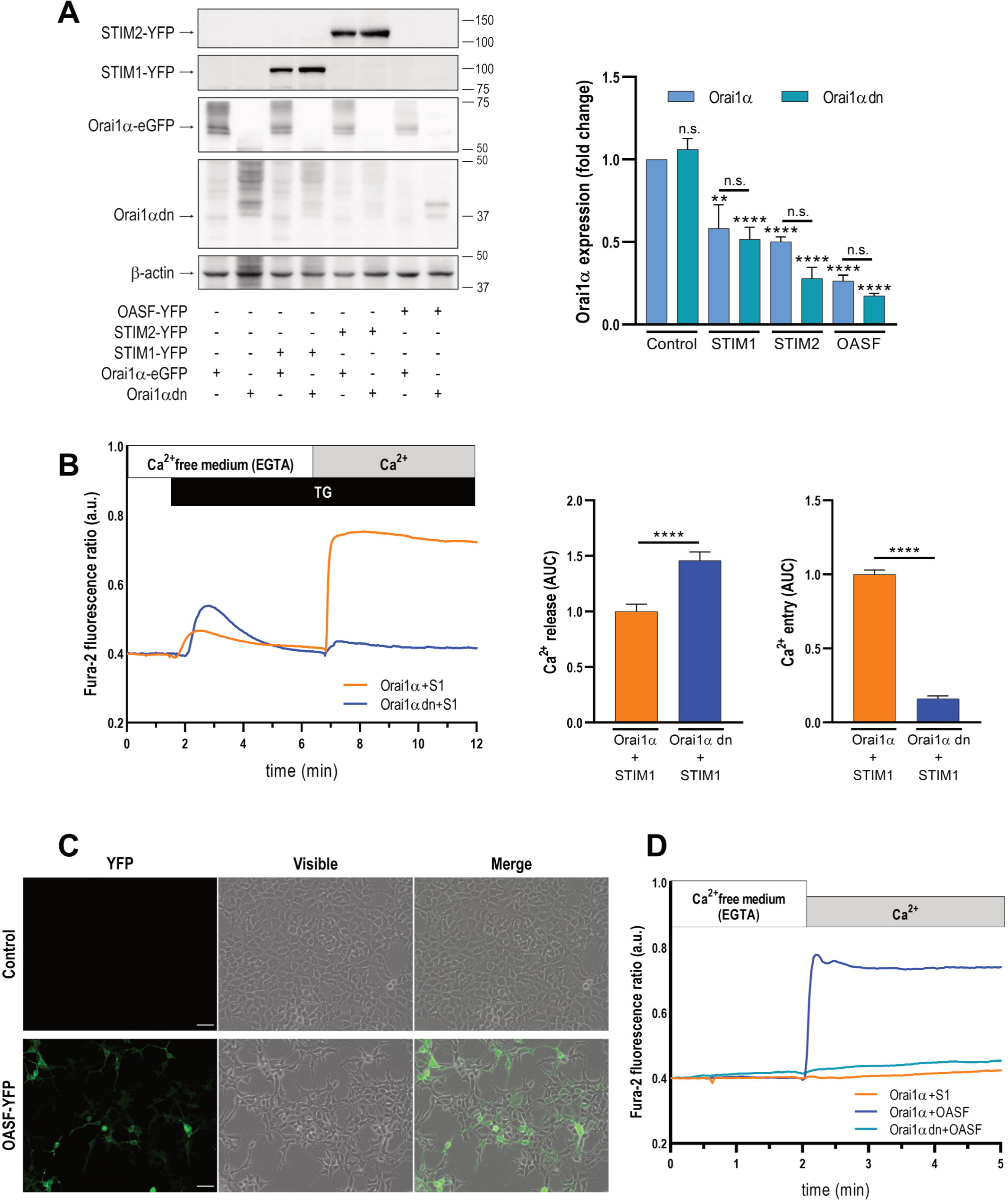
Degradation of Orai1α and Orai1β by STIMs does not require Ca^2+^ influx through the channel. (**A**) STIM1,2-DKO HEK-293 cells were co-transfected with CMV-driven Orai1α-eGFP or a dominant negative Orai1α mutant (Orai1αdn), in combination with either STIM1-YFP, STIM2-YFP or the STIM1 OASF fragment, as indicated. Forty-eight hours later cells were lysed and cell lysates were analyzed by SDS-PAGE and Western blot analysis using anti-STIM1, anti-STIM2 or anti-Orai1 C-terminal antibody. Membranes were reprobed with anti β-actin antibody for protein loading control. Molecular masses indicated on the right were determined using molecular-mass markers run in the same gel. These results are representative of 6 separate experiments. Bar graphs show quantification of Orai1α and Orai1αdn expression under the different experimental conditions normalized to the β-actin expression. Data are represented as mean[±[SEM and expressed as fold change (experimental/control). Data were statistically analyzed using Kruskal–Wallis test with multiple comparisons (Dunn’s test). \*\**P* < 0.01 and \*\*\*\**P* < 0.0001 as compared to the expression of Orai1α or Orai1αdn in the absence of STIM proteins and fragments. (B) STIM1,2-DKO HEK-293 cells were co-transfected with CMV-driven Orai1α or a dominant negative Orai1α mutant (Orai1αdn), in combination with STIM1, as indicated. Forty-eight hours later cells were loaded with fura-2 and suspended in a Ca^2+^-free HBS (100 µM EGTA added) and then stimulated with TG (2 µM) followed by the addition of extracellular Ca^2+^ (final concentration 1.8 mM) to initiate Ca^2+^ influx. Bar graphs represent quantification of TG-evoked Ca^2+^ release and entry determined as described in materials and methods. Data are presented as mean ± SEM and expressed as fold change over control (cells expressing Orai1α and STIM1). Data were statistically analyzed using Mann-Whitney U test. \*\*\*\**P* < 0.0001 as compared to cells expressing Orai1α and STIM1. (**C**) STIM1,2-DKO HEK-293 cells were co-transfected with CMV-driven STIM1 OASF fragment. Forty-eight hours later GFP fluorescence was detected using an LSM900 confocal microscope. The images show representative confocal images of the STIM1 OASF region. The scale bar represents 100 μm. (**D**) STIM1,2-DKO HEK-293 cells were co-transfected with CMV-driven Orai1α or a dominant negative Orai1α mutant (Orai1αdn), in combination with STIM1 or the STIM1 OASF fragment, as indicated. Forty-eight hours later cells were loaded with fura-2 and suspended in a Ca^2+^-free HBS (100 µM EGTA added). Ca^2+^ was added to the extracellular medium at a final concentration of 1.8 mM to initiate Ca^2+^ influx.

Next, we further explored whether STIM1-Orai1α interaction is required for Orai1 degradation and, hence, we have tested the protein content of Orai1α in the presence of STIM1 or the STIM1-OASF (Orai1 activating small fragment) region (aa 233-474) [6]. As expected, co-expression of Orai1α and OASF resulted in a robust activation of constitutive Ca^2+^ entry independently of Ca^2+^ store depletion as compared to cells expressing Orai1α and STIM1 (Fig. 11C-D). The constitutive Ca^2+^ influx observed in OASF-expressing cells was blocked when co-expressed with the Orai1dn mutant (Fig. 11D). As depicted in Fig. 11A, the Orai1α protein content was reduced either when co-expressed with STIM1 or the STIM1-OASF region. We observed that the Orai1α protein level was even smaller when co-expressed with the OASF region (Fig. 11A), which we attribute to a greater interaction of Orai1α with OASF than with full-STIM1 at rest. These findings indicate that interaction of Orai1α with the OASF region is sufficient to promote Orai1α degradation. Co-expression of OASF with Orai1dn also induced a drop in the Orai1dn protein content. The degradation of Orai1dn was similar to that observed with Orai1α, which further confirms that Ca^2+^ influx through the channel is not required for this event.

### Orai1α and Orai1β promote STIM1 turnover

Since STIM proteins promote Orai1 recycling we wonder whether, reciprocally, Orai1 facilitates the turnover of STIM proteins. To address this question, we expressed STIM1 or STIM2 in STIM1/2 DKO cells alone or in combination with Orai1α and Orai1β and analyzed the protein content of the STIM proteins. Our results indicate that when STIM1 was co-expressed with Orai1α or Orai1β its protein content was significantly reduced (Fig. 12A; *P*<0.0001). By contrast, the protein content of STIM2 was not found to be affected by expression of the Orai1 variants (Fig 12B). These findings suggest that STIM1 turnover is promoted by Orai1α and Orai1β while STIM2 escapes from this regulatory mechanism.

**Fig. 12.**
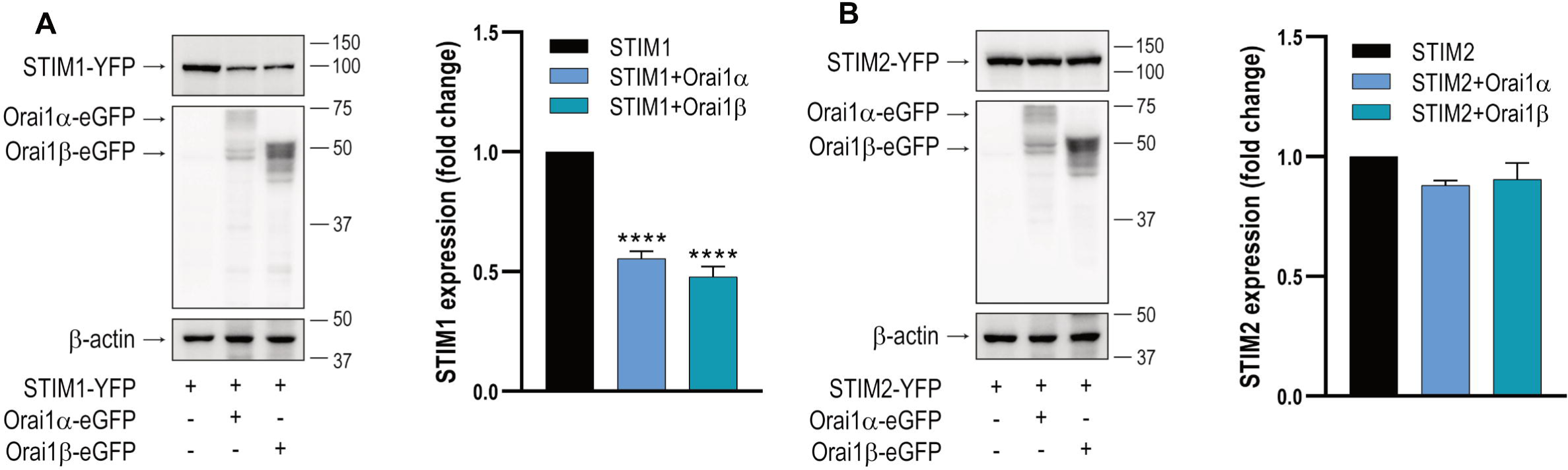
STIM1 stability is modulated by Orai1α and Orai1β. STIM1,2-DKO HEK-293 cells were co-transfected with CMV-driven STIM1-YFP (**A**) or STIM2-YFP (**B**) in combination with Orai1α-eGFP or Orai1β-eGFP, with a relative Orai1:STIM expression ratio of 1:2. Forty-eight hours later cells were lysed and cell lysates were analyzed by SDS-PAGE and Western blot analysis using anti-STIM1 or anti-STIM2 antibody. Membranes were probed with anti-Orai1 C-terminal antibody or anti β-actin antibody. Molecular masses indicated on the right were determined using molecular-mass markers run in the same gel. These results are representative of 6 separate experiments. Bar graphs represent the quantification of STIM1 and STIM2 expression under the different experimental conditions normalized to the β-actin expression. Data are represented as mean[±[SEM and expressed as fold increase (experimental/control). Data were statistically analyzed using Kruskal–Wallis test with multiple comparisons (Dunn’s test). \*\*\*\**P* < 0.0001 as compared to the STIM1 expression in the absence of Orai1 variants.

## Discussion

It is well established that mammalian STIM proteins, STIM1 and STIM2, differ in their properties and biological functions. STIM1 is the main activator of Orai/CRAC channels while STIM2, with lower affinity to Ca^2+^, has been proposed as a slower and weaker Orai activator that stabilizes basal cytosolic and ER Ca^2+^ levels [5]. In agreement with the previous observations [5, 35] here we show that STIM1 is a stronger activator that STIM2 triggering Orai1α-dependent Ca^2+^ signals. By contrast, we provide for the first time evidence supporting that STIM1 and STIM2 exhibit a similar strength activating Ca^2+^ signals generated by Orai1β. Native CRAC channels have been reported to consist of heteromers of the three Orai isoforms [26], and previous studies have shown that STIM2 exhibits basal interaction with Orai1 and Orai2, a process that might regulate basal cytosolic and ER Ca^2+^ levels and SOCE upon stimulation with low agonist concentrations [1]. Concerning the Orai1 variants, our FRET analysis demonstrates a significantly enhanced basal interaction between STIM2 and Orai1α, as compared to Orai1β, a difference that has not been observed between STIM1 and the Orai1 forms. This observation might suggest that Orai1α would play a major role in the modulation of resting cytosolic and ER Ca^2+^ levels.

The canonical role of STIM proteins is to communicate the filling state of the intracellular Ca^2+^ stores to the Orai channels in the PM to mediate SOCE. Modulation of Ca^2+^ entry through Orai channels is essential to generate physiological Ca^2+^ signals and maintain intracellular Ca^2+^ homeostasis. A variety of regulatory mechanisms have been reported, including intrinsic mechanisms such as fast Ca^2+^-dependent inactivation (FCDI) of Orai channels [36] and, based on the different sensitivity of the three Orai isoforms to FCDI, fine-tuning of Ca^2+^ entry by Orai heteromerization [26]. Other SOCE regulatory mechanisms require the participation of regulatory proteins and/or organelles such as slow Ca^2+^-dependent inactivation [37, 38]. Here we provide evidence for a role of STIM1 and STIM2 in the turnover of Orai1α and Orai1β that might serve as an additional feedback modulator of SOCE. We base this statement in different findings: First, in STIM1/2 DKO cells, expression of STIM1 and STIM2 significantly attenuated the protein content and PM location of Orai1α and Orai1β. This difference cannot be attributed to interference between the expression plasmids as we did not detect changes in the expression of the Orai1 variants upon co-expression of Orai1α with Orai1β or with other cellular proteins using the same CMV-driven plasmids and experimental procedure [21]. Concerning the PM expression of the Orai1 variants, the expression of STIM proteins did not alter the increase in the PM location of Orai1 observed upon Ca^2+^ store depletion relative to control, which confirms that STIM proteins modulate the protein content of Orai1 but not its functionality. Our previous studies using Orai1 knockout HEK-293 cells have reported that while at resting conditions there is a pool of both Orai1 variants in the PM, upon Ca^2+^ store depletion there is a detectable PM enrichment of Orai1α while Orai1β behaves as a defective protein that requires the presence of Orai1α to accumulate in the PM [21]. In the present study we have detected an increase in the surface expression of transiently expressed Orai1β after stimulation with TG in the absence of cell transfection with Orai1α, which we attribute to the presence of native Orai1α in these cells. Second, the extent of the drop in Orai1 variant protein content depends on the amount of STIM expressed as observed when DKO cells were co-transfected with STIM and Orai1 variants expression plasmids at 1:1 and 1:2 ratios. Third, the STIM protein content modulates the expression of native Orai1α and Orai1β as demonstrated when WT HEK-293 cells were transfected with STIM1 or STIM2 expression plasmid or, alternatively, with shRNA/siRNA specific for each STIM protein. Forth, STIM-induced Orai1α and Orai1β turnover is likely to involve interaction between both proteins as demonstrated by the enhanced drop in the Orai1 protein level induced by the constitutively active mutants STIM1-D76A and STIM2-D80A or the STIM1-OASF region. To our knowledge this is the first evidence for a role of STIM proteins in the regulation of Orai1 variant content, and, in this regulatory function, the role of both STIM proteins is indistinguishable.

The content of a given protein depends on the rate of synthesis and degradation. By using CHX, which blocks protein synthesis by impairing translation elongation [39] we have found that STIM proteins do not alter the synthesis of the Orai1 variants, as the percentage of inhibition of their expression in the presence of STIMs is similar when protein synthesis is allowed or impaired. Similarly, inhibition of ubiquitin-proteasomal degradation has negligible effects, if any, on the Orai1 protein content. By contrast, impairment of the lysosomal proton gradient using bafilomycin A1 completely reversed the effect mediated by expression of STIM1 and STIM2, which demonstrates that STIM proteins modulate the protein level of Orai1α and Orai1β by endo-lysosomal degradation. Our findings are consistent with a previous study that revealed the role of lysosomal degradation in Orai1 protein turnover [40]. Furthermore, we have confirmed that the STIM1 OASF region mimics the degradation of Orai1 caused by STIM1, which indicates that OASF is sufficient for this process and strongly suggests that Orai1 degradation requires OASF-Orai1 interaction. Nevertheless, Orai1 degradation does not depend on Ca^2+^ influx through the channel, as expression of STIM proteins induces degradation of Orai1α bearing the pore-dead E106Q mutation.

We have found that the protein content of STIM1 is modulated by the expression of the Orai1 variants, which might suggest that both proteins recycle simultaneously. Interestingly, we have not detected any significant drop in the STIM2 protein content after co-expression with the Orai1 variants, thus, indicating that STIM2 recycles by a mechanism that is independent of Orai1. As for Orai1, STIM1 and STIM2 recycling has been previously reported to be mediated by autophagy/lysosomal degradation and is independent of the proteasome [41, 42]. However, the half-life of STIM2 (>24h) [43] has been reported to be significantly greater than that of STIM1 (19h) [44]. The results presented here, demonstrating that STIM2 recycling is independent of Orai1 interaction, provides an explanation for the greater stability of STIM2 and are consistent with a role for STIM2 in the maintenance of basal Ca^2+^ levels.

Therefore, our results provide evidence that support two important aspects of the regulation of Orai1 variants function. On the one hand, while STIM1 is a stronger activator of Orai1α than STIM2, both STIM proteins exhibit similar strength activating Orai1β-dependent Ca^2+^ signals. Nevertheless, we have found that there is a significantly greater basal interaction of STIM2 with Orai1α that might suggest that Orai1α plays a major role in the regulation of basal cytosolic and ER Ca^2+^ levels. Furthermore, our results provide evidence supporting that STIM proteins play a relevant role in Orai1α and Orai1β recycling, which might serve as an additional feedback regulatory mechanism of SOCE as the extent of Ca^2+^ influx through Orai1α and Orai1β is expected to depend not only on the intrinsic properties of the channels itself, such as Ca^2+^-dependent inactivation, but also on the number of available channels and its PM location.

## Acknowledgments.

We thank Sandra Alvarado for assistance in some experiments and María Barriga for technical assistance.

## Author contributions

Conceptualization of manuscript, I.J, T.S., J.A.R.; data curation, I.J., J.A.R.; formal analysis, J.N-F., I.J.; funding acquisition, T.S., J.A.R.; investigation, J.N-F., I.J., A.M-D., V.J-V.; methodology, J.N-F., I.J., A.M-D., J.J.L.; project administration, J.A.R.; resources, J.A.R.; Supervision, I.J, T.S., JAR.; Validation, I.J, J.A.R.; Writing-original draft, I.J, J.A.R.; Writing-review and editing, G.M.S., J.J.L., I.J, T.S., J.A.R.

## Funding.

This research was funded by PID2022-136279NB-C21 and PID2022-136279NB-C22 funded by MCIN/AEI/10.13039/501100011033 and by “ERDF A way of making Europe”, and Junta de Extremadura-Fondo Europeo de Desarrollo Regional (FEDER; Grant IB20007 and GR18061) to JAR and TS. JN-F and AM are supported by a contract from AEI/Spanish Ministry of Science and Innovation and INVESTIGO, respectively.

## Data and material availability

All data are available in the main text or the supplementary materials.

## Declarations

### Conflict of interest

The authors declare no competing interests.

### Ethical approval

Experimental procedures were approved by the local ethical committee (University of Extremadura and Extremadura Health Service).

### Consent for publication

We, all the authors, give our consent for publication.

## Notes

### Competing Interest Statement

The authors have declared no competing interest.

