## Supplementary Fig 1 for "Feedback modulation of Orai1α and Orai1β protein content mediated by STIM proteins"

### UNCROPPED GELS

#### Figure 1F

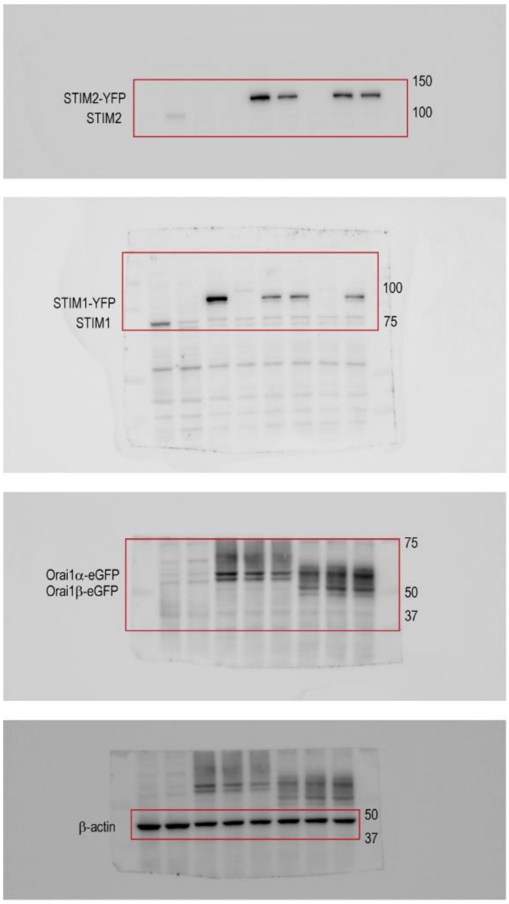


#### Figure 3F

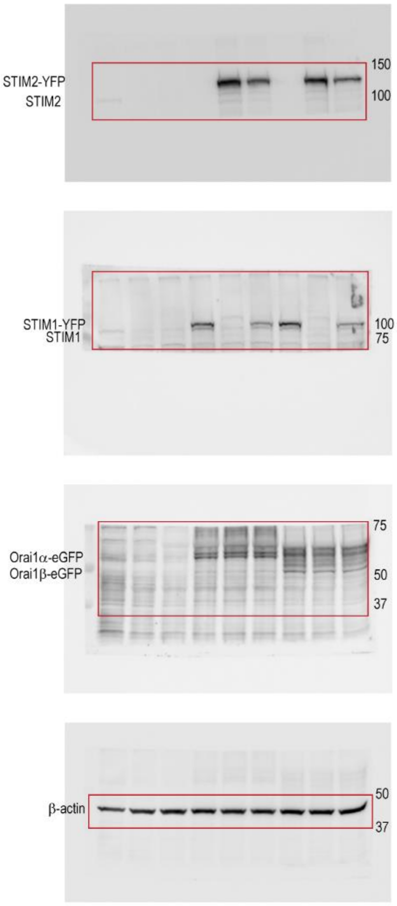


#### Figure 6A and 6B

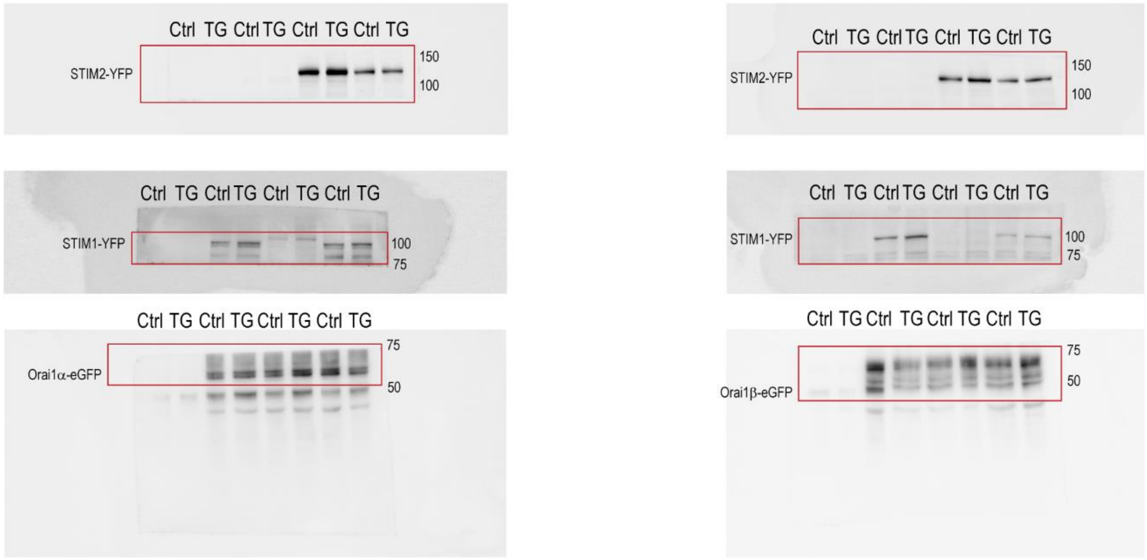


**Supp. Figure continue**

### Figure 7A and 7B

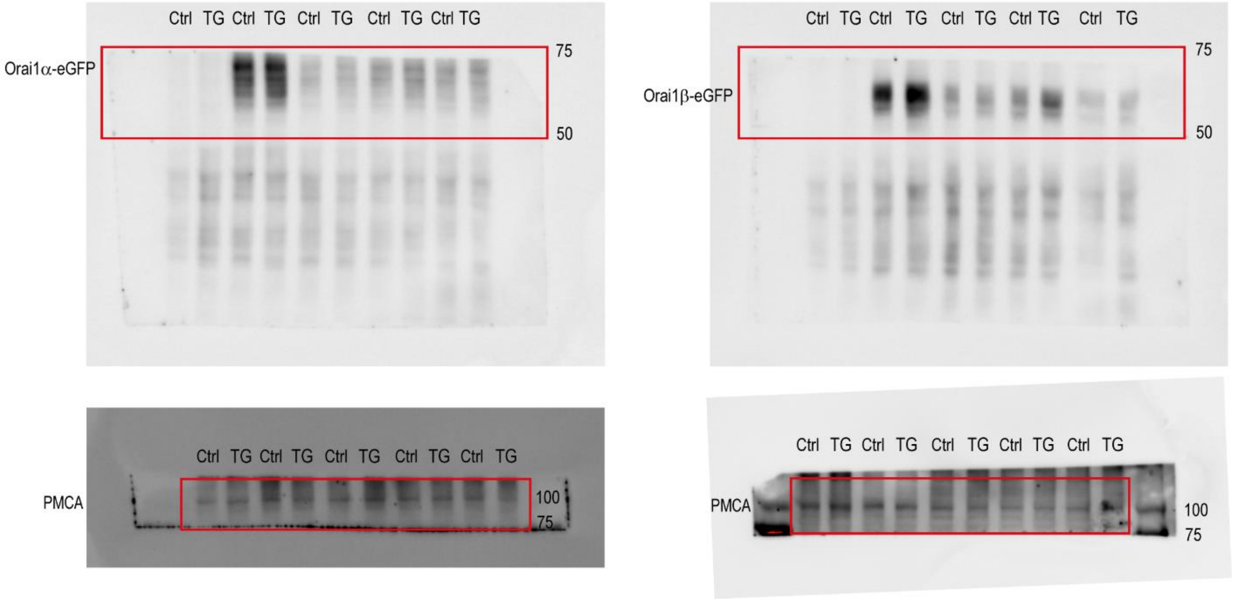


### Figure 8A and 8B

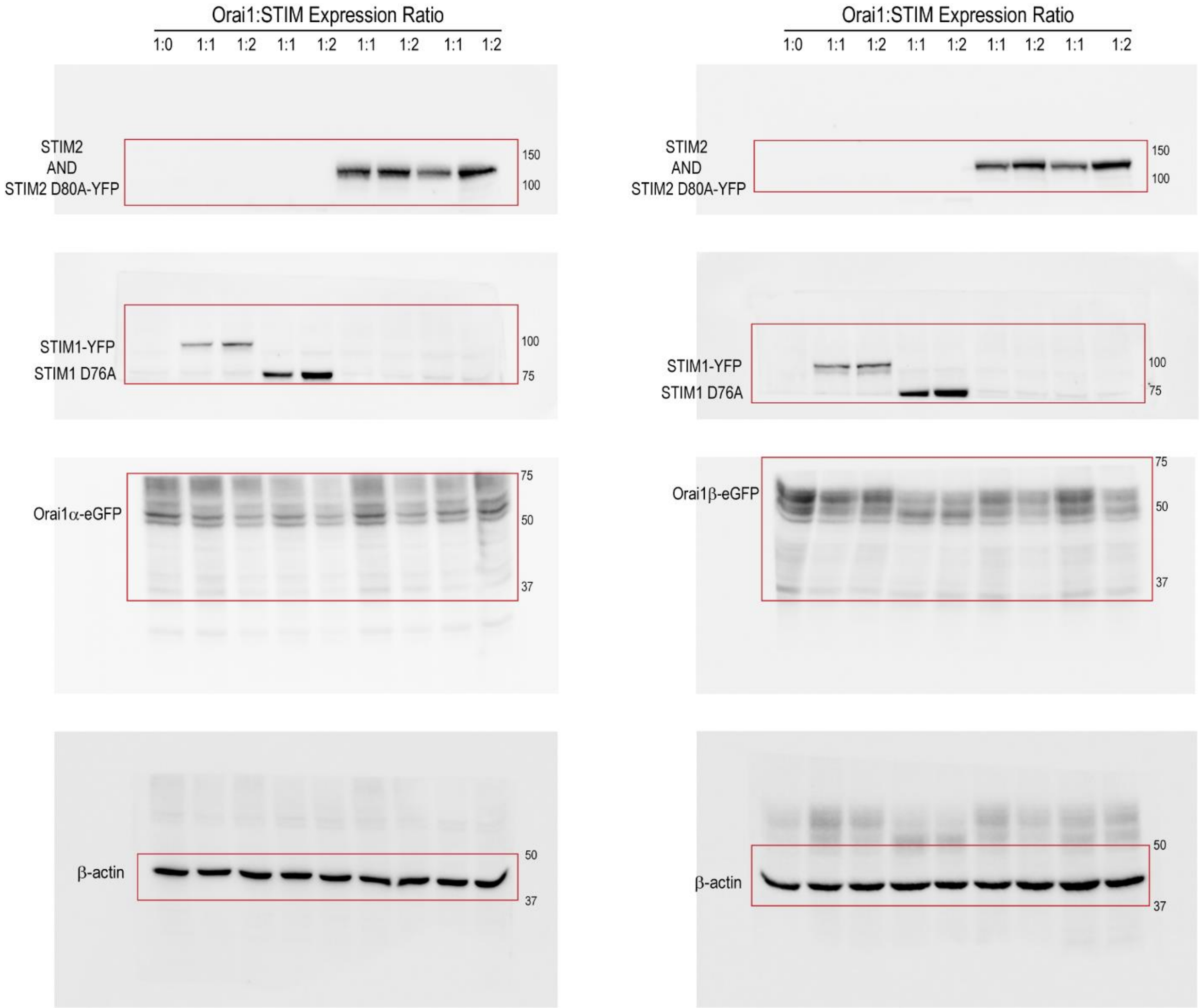


**Supp. Figure continue**

### Figure 9A

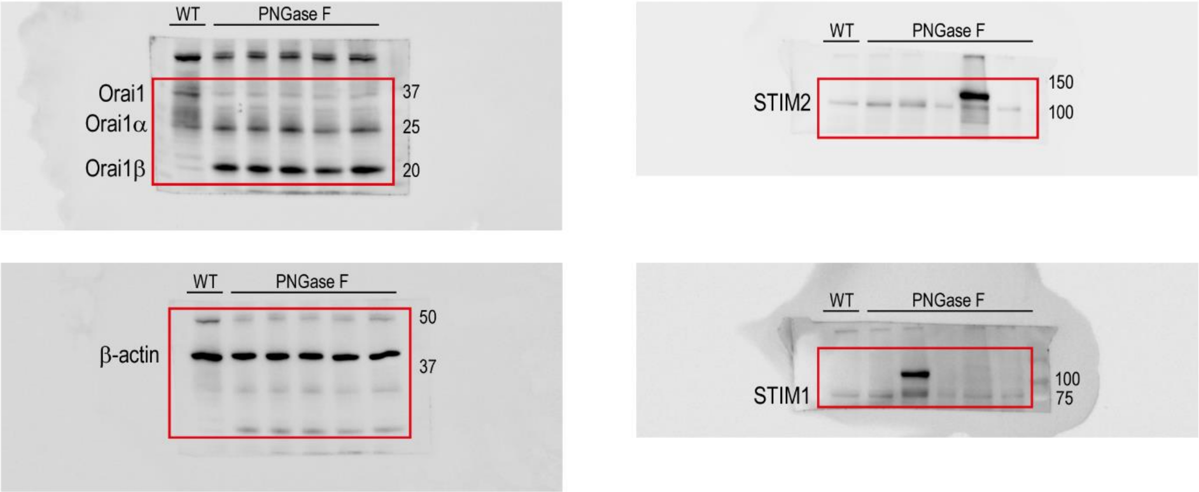


### Figure 10A and 10B

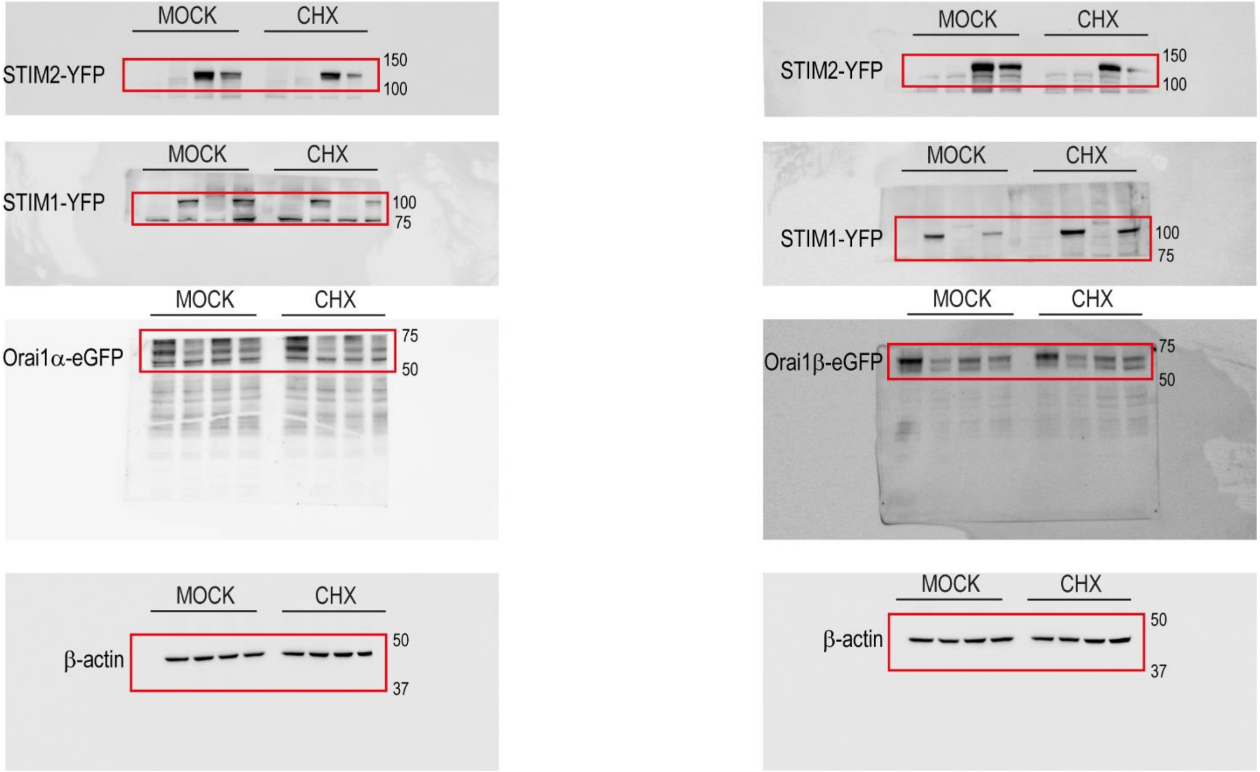


**Supp. Figure continue**

### Figure 10C and 10D

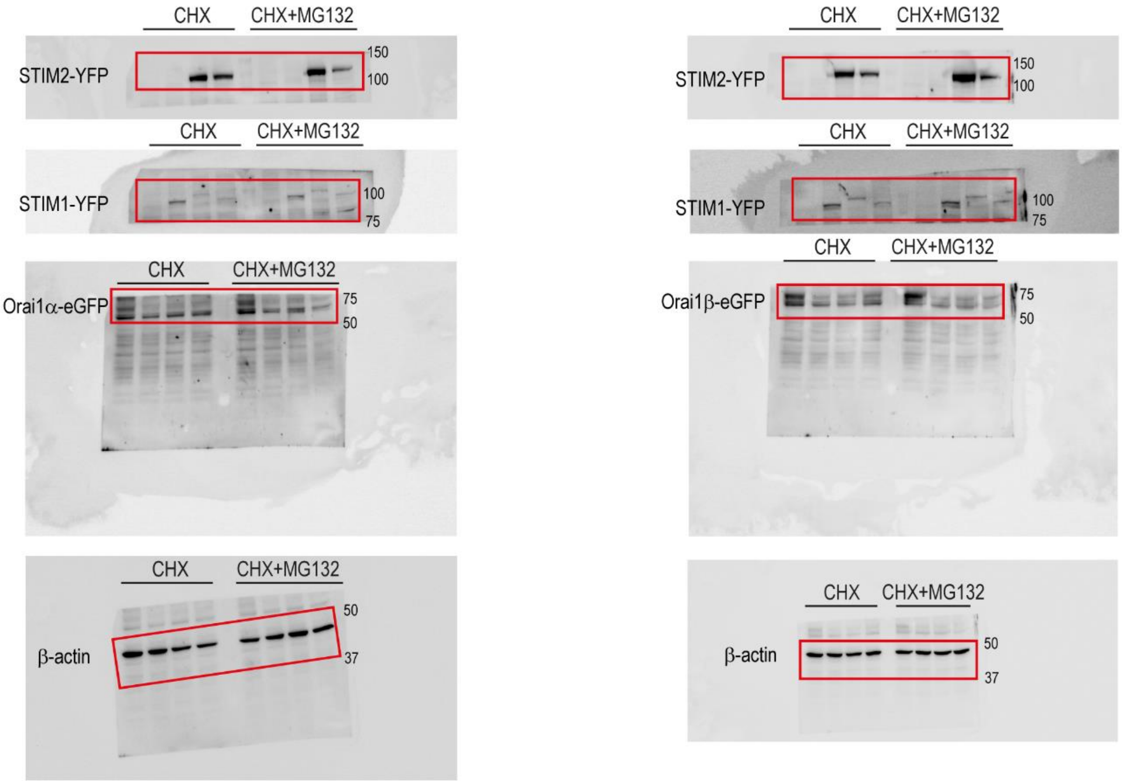


### Figure 10E and 10F

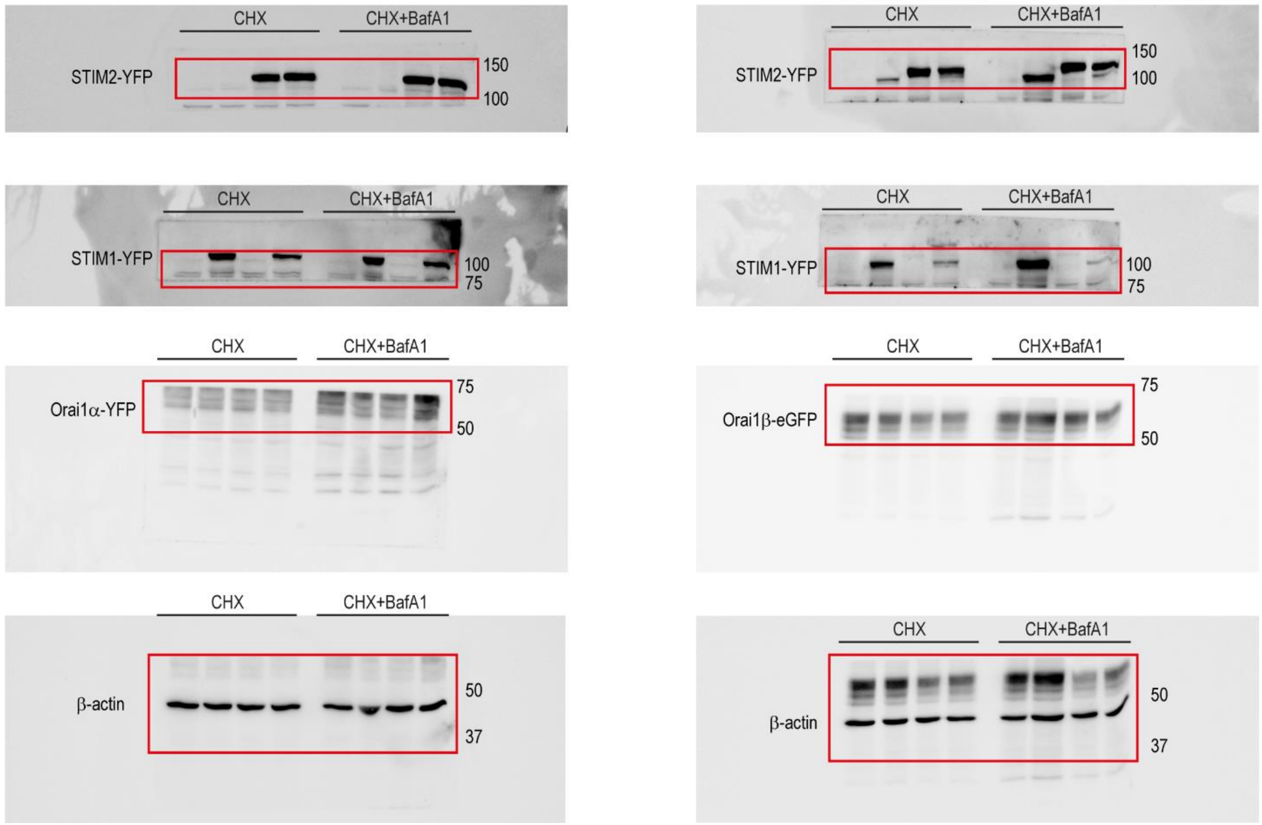


**Supp. Figure continue**

### Figure 11A

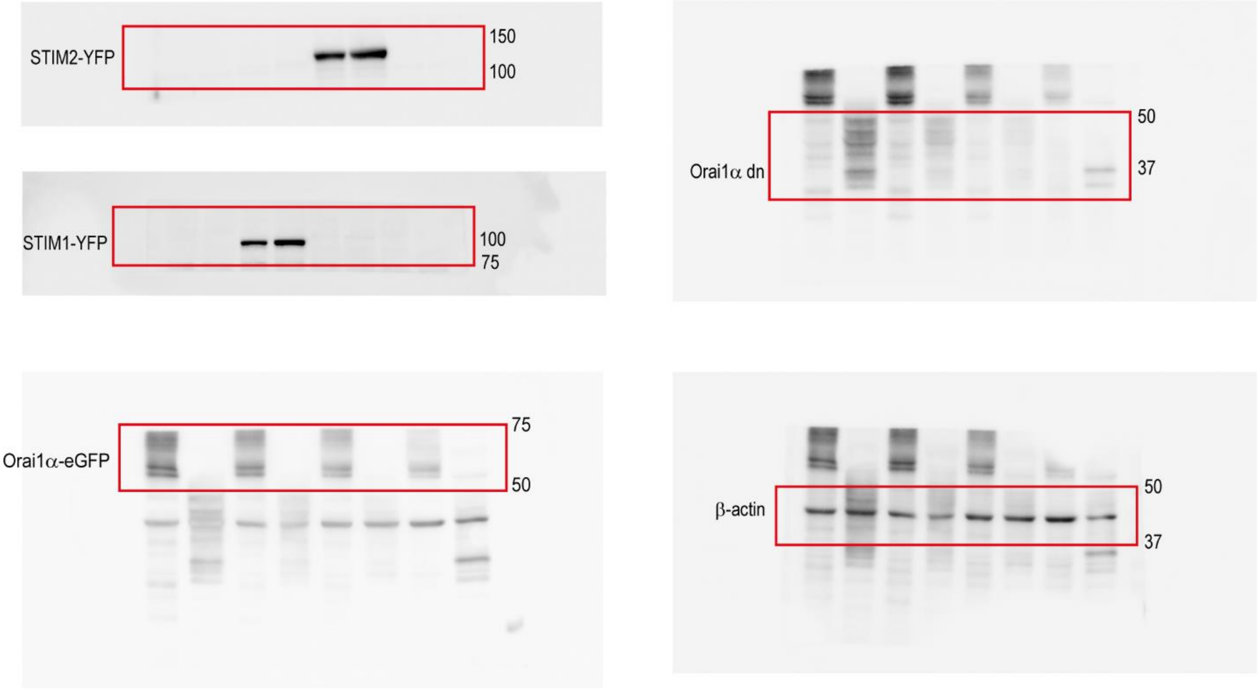


### Figure 12A and 12B

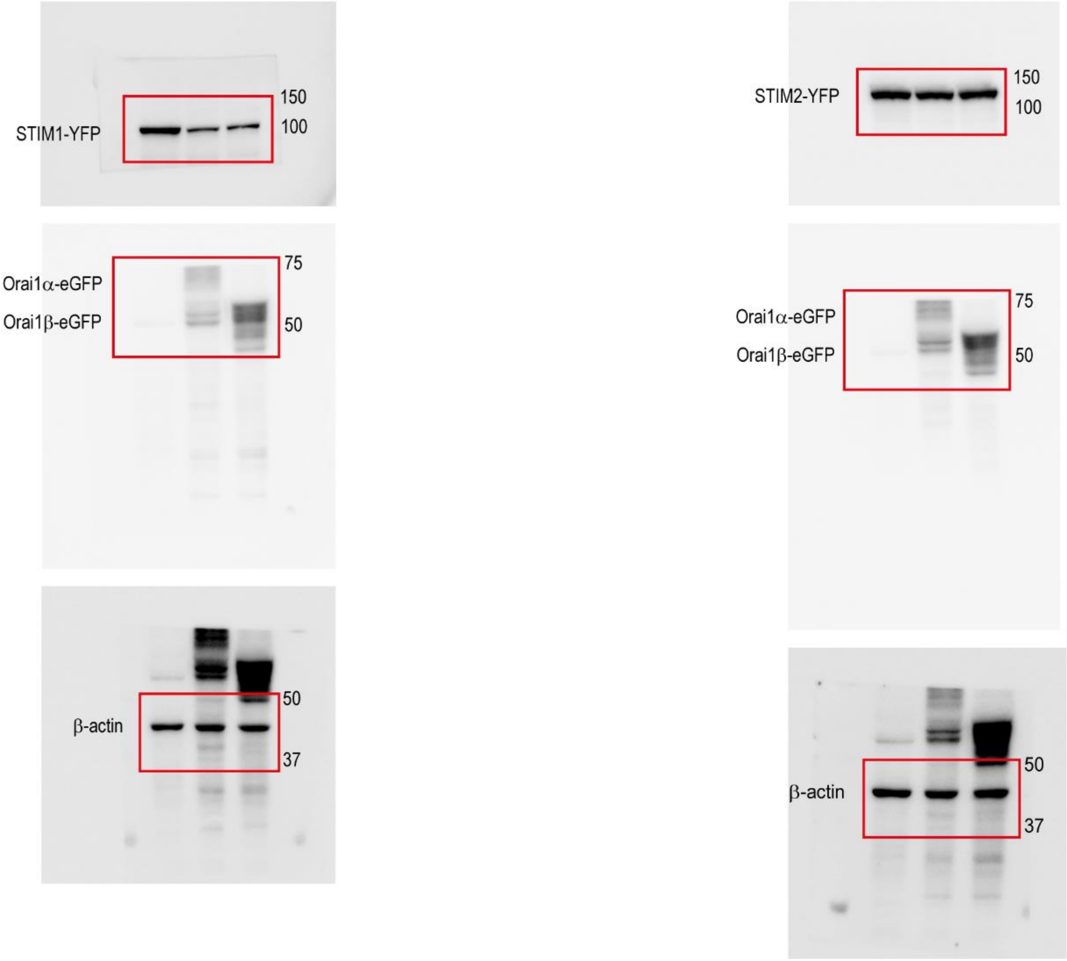
